# Intrathecal (G_4_C_2_)_149_ delivery in C9orf72-deficient mice yields mild motor dysfunction and ALS/FTD pathological hallmarks

**DOI:** 10.64898/2026.02.09.704060

**Authors:** Katelyn A. Russell, Amelia A. Shahrabi, Suleyman C. Akerman, Matthew D. Byrne, Jeffrey D. Rothstein, Davide Trotti, Brigid K. Jensen, Aaron R. Haeusler

**Author notes:** Correspondence: Aaron R. Haeusler, Ph.D. Jefferson Weinberg ALS Center, Vickie and Jack Farber Institute for Neuroscience Department of Neuroscience, Thomas Jefferson University Philadelphia, PA 19107, USA.

## Abstract

A repeat expansion in *C9orf72* is the most common genetic cause of amyotrophic lateral sclerosis (ALS) and frontotemporal dementia (FTD), yet existing mouse models incompletely engage spinal regions implicated in disease. Here, an adeno-associated virus encoding (G_4_C_2_)_149_ repeats was delivered via neonatal intrathecal injection, achieving widespread CNS expression with robust spinal cord targeting. This approach was applied to mice with graded loss of endogenous *C9orf72* to interrogate both gain- and loss-of-function mechanisms. Longitudinal motor, behavioral, and pathological analyses revealed that repeat expression primarily drives mild, progressive muscle weakness, whereas coordination deficits were largely genotype dependent. Subtle gait abnormalities and hyperactivity were also observed. Within spinal motor regions, repeat-expressing mice exhibited dipeptide repeat protein accumulation, reduced NeuN-positive area, glial activation, and sparse phosphorylated TDP-43 pathology. Cross-domain correlations further linked repeat expression, spinal pathology, and motor dysfunction. Collectively, these findings establish that CNS-wide repeat expression combined with reduced *C9orf72* produces a coherent, mild ALS/FTD model.

## INTRODUCTION

Amyotrophic lateral sclerosis (ALS) and frontotemporal dementia (FTD) are fatal neurodegenerative diseases historically considered distinct but now recognized as a clinical^1–5^, pathological^6–8^, and genetic^9^ disease spectrum. ALS is characterized by progressive upper and lower motor neuron degeneration, with heterogeneous presentations ranging from muscle weakness and atrophy to spasticity, hyperreflexia, or bulbar symptoms^10–13^. FTD, an early-onset dementia affecting the frontal and temporal lobes, presents with cognitive and behavioral deficits, including diminished executive function and impaired judgment^14,15^. Despite these differing primary features, ∼15% of sporadic ALS patients develop FTD^2^, while 12.5–14% of FTD patients develop motor neuron disease^1,16^; cognitive impairment occurs in over 50% of sporadic^2^ and up to 60% of familial ALS^3^, and 27–36% of FTD patients exhibit motor dysfunction^1,16^.

Pathologically, aggregation of TDP-43 is a shared hallmark observed in over 97% of ALS cases and approximately 50% of FTD cases^6–9^. Disease-associated TDP-43 features nuclear depletion and/or cytoplasmic mislocalization, causing loss of normal nuclear function and toxic gain-of-function effects^17–19^, linking these disorders through shared molecular pathology.

A hexanucleotide (G_4_C_2_)*_n_* repeat expansion in the *C9orf72* gene (C9-NRE) is the most frequent known genetic driver of the ALS/FTD spectrum. Repeat lengths range from fewer than 24 in unaffected individuals to hundreds or thousands in patients, with variability across tissues and cell types^20–24^. Accounting for ∼40% of familial ALS, ∼25% of familial FTD, ∼7% of sporadic ALS cases, and ∼7% of sporadic FTD cases^4,20,21,25–28^, the C9-NRE provides a shared molecular framework for dissecting disease mechanisms.

Three non-mutually exclusive mechanisms have been proposed to underlie C9-NRE toxicity: loss of C9orf72 protein function, RNA-mediated toxicity through repeat-containing transcripts, and gain-of-function toxicity from dipeptide repeat proteins (DPRs) generated by repeat-associated non-ATG translation^7,24^. Evidence from patient tissue and experimental models supports contributions from each mechanism, suggesting ALS/FTD pathogenesis may arise from their convergence rather than a single dominant pathway^7,24,29^.

Additionally, neuroinflammation is a central feature of ALS/FTD, including C9-NRE-associated disease. PET imaging and postmortem analyses reveal microglial proliferation and activation in regions of neuronal loss, with inflammatory burden correlating to functional decline^30–38^. Proteomic network analyses show enrichment of glial proteins in ALS/FTD molecular modules, with C9-NRE-specific changes mapping to microglial and astrocytic networks^39^. Supporting these observations, *C9orf72* loss-of-function mouse models exhibit immune dysregulation and age-related neuroinflammation^40–42^. While these models lack overt motor neuron degeneration, indicating *C9orf72* haploinsufficiency alone is insufficient to drive ALS pathology, these findings suggest inflammation may interact with repeat-dependent gain-of-function mechanisms to influence disease progression.

To investigate repeat-dependent toxicity, multiple mouse models have been developed using genetic and viral strategies. Adeno-associated virus (AAV)-mediated intracerebroventricular (ICV) injection of expanded (G_4_C_2_)*_n_*repeats is widely used to study C9-NRE gain-of-function. Models expressing (G_4_C_2_)_66_ or (G_4_C_2_)_149_ repeats recapitulate key pathological features, including RNA foci, DPR accumulation, and gliosis, alongside variable neurodegeneration, pTDP-43 pathology, and behavioral phenotypes^43–46^. Although phenotypes vary and motor deficits are often modest, these models provide critical insights into C9-NRE molecular and cellular pathology.

Combinatorial models^29^ have revealed synergistic interactions, incorporating both C9-NRE expression and reduced *C9orf72* function. Augmenting loss-of-function mice with C9-NRE expression accelerated DPR accumulation, enhanced neuronal loss, and exacerbated cognitive deficits. Importantly, combinatorial models also developed phenotypes not observed in the parent models^40,43,47^, including increased neuroinflammation, reduced autophagy, and age-dependent deficits in motor coordination. Collectively, these studies highlight the importance of modeling both *C9orf72* gain- and loss-of-function mechanisms.

Despite these advances, existing C9-NRE mouse models do not robustly recapitulate spinal pathology or motor dysfunction, defining features of ALS. Here, we evaluated whether CNS-wide C9-NRE expression, including the spinal cord, combined with graded *C9orf72* loss is sufficient to drive disease-relevant motor phenotypes. To this end, an AAV encoding (G_4_C_2_)_149_ repeats was intrathecally injected into neonatal mice with intact, partial, or complete *C9orf72* loss. This approach achieved widespread C9-NRE expression, including robust engagement of spinal motor regions underrepresented in existing models. Longitudinal motor, behavioral, and pathological analyses revealed that CNS-wide C9-NRE expression, alongside reduced *C9orf72* function, induces progressive muscle weakness, impaired coordination, subtle gait and cognitive changes, and hallmark pathology including DPR accumulation, reduced NeuN-positive signal, glial activation, and sparse pTDP-43 inclusions. Together, these findings establish a model integrating gain- and loss-of-function mechanisms that recapitulates mild yet disease-relevant motor phenotypes alongside convergent pathological features of ALS/FTD, providing a platform for mechanistic studies and therapeutic development targeting motor and spinal cord-associated phenotypes.

## METHODS

### Experimental models

All animal procedures were performed in accordance with protocols reviewed and approved by the Institutional Animal Care and Use Committee (IACUC) of Thomas Jefferson University.

### Mice

*C9orf72* knockout mice were purchased from The Jackson Laboratory (C57BL/6J-C9orf72em5Lutzy/J; strain #: 027068) and bred to produce a colony of wildtype, heterozygous, and homozygous *C9orf72* knockout mice. Mice were housed in standard cages in a light-, humidity-, and temperature-controlled facility and provided food and water ad libitum. Breeding cages were supplied with higher fat (6% fat) LabDiet 5K52 food to support reproduction and lactation. Upon weaning, mice were switched to regular (4.5% fat) LabDiet 5010.

For all Motor Performance assessments and Cognitive and Behavior Measures, mice were brought to the testing room and allowed to acclimate for roughly 30 minutes.

### AAV Injections

Frozen aliquots of (G_4_C_2_)_149_ repeat-containing AAV were provided by Dr. Leonard Petrucelli and his lab at Mayo Clinic (Jacksonville, FL). Viruses, produced and processed as previously described^44^, were stored at −80°C until use. To prepare for injections, aliquots of AAV were thawed on ice then spun down. In a sterile hood, AAV was diluted in sterile PBS with 0.05% Trypan Blue, then kept on ice. For intrathecal injections, based on neonatal mouse AAV intrathecal injection and neurodegenerative literature^48–54^, each mouse received a dose of 8.25×10^10 genomes in a total volume of 15μL. For intracerebroventricular (ICV) injections, based on previous C9-NRE ICV mouse models^43^, each mouse received 4.00×10^10 genomes total, split between both lateral ventricles (2μL injected into each ventricle). All injections took place at postnatal day 1 (P1).

As previously described^48–50,52^, pups receiving intrathecal injections were cryo-anesthetized. At P1 the spinal cord is visible through the skin as a white stripe down the pup’s back. A 34G needle (Hamilton 207434-10) was inserted at the mid-line in the upper lumbar vertebral column, about 5 mm from the base of the tail, and either AAV or control solution was injected. Successful injections were indicated by the appearance of Trypan blue in the spinal cord, cerebellum, and cranial sutures, as well as no leakage of the injectant when the needle was withdrawn. Only pups with successful injections were included in the study. After the injection, the pup was warmed by hand until limb movement was detected, then returned to the mother.

Pups receiving ICV injections were cryo-anesthetized. As previously described^29,43–45^, each injection site was identified, two-fifths of the distance from the lambda suture to the eye, and marked with a laboratory pen. While gently stabilizing the pup on its side, the needle was inserted perpendicular to the skull surface, roughly 3 mm deep or when the resistance on the needle decreased, indicating the tip had reached the lateral ventricle, and either AAV or dye control was slowly injected. An injection was considered successful when the Trypan blue (visible through the skin and skull) filled the lateral ventricle yielding a football-like or oval shape, and when there was no leakage of injectant when the needle was removed. Both injections, one into each lateral ventricle, had to be successful for the pup to be considered for the study. After both lateral ventricles were injected, the pup was warmed by hand until limb movement was detected, then returned to the mother.

GFP and *C9orf72* viruses were AAV2/9. The C9-NRE virus contained a (G_4_C_2_)_149_ repeat expansion flanked by regions of the *C9orf72* gene (119 base pair 5’ and 100 base pairs 3’), under a chicken beta-actin promoter^44^. The GFP reporter virus was comparable to the C9-NRE, but contained a fluorescent tag to track expression patterns instead of the (G_4_C_2_)_149_ repeat expansion^43^. Control injections consisted of sterile PBS and 0.05% Trypan Blue. Additionally, some mice received no injection but were not found to be significantly different from the Control injection mice. Therefore, Control injection and non-injected groups were combined as Control for analysis.

### Motor Performance

#### Electrophysiology

Mice were anesthetized with ∼2% isoflurane. Compound muscle action potentials (CMAPs) were recorded from the sciatic nerve-footpad system using the Neurosoft Neuro-MEP micro and Neurosoft Neuro-MEPomega (version 3.7.3.10; Neurosoft, Ivanovo, Russia) and 28G stainless steel needle electrodes (Technomed Medical Accessories, the Netherlands). Stimulation electrodes were placed at two sites along the left sciatic nerve, the proximal site at the sciatic notch and the distal site at the ankle, near the Achilles tendon. Recording electrodes were inserted at the plantar muscles of the footpad. Stimulation voltages were adjusted to elicit maximal CMAP response and consisted of short pulses (<0.2 ms). CMAP amplitude was measured peak-to-peak^55–57^.

#### Grip Strength

Grip was assessed using the Chatillon Ametek Grip Strength Meter (Columbus Instruments) with the triangle grip bar. Mice were guided up to the bar, supported by the tail and forelimbs, positioned horizontally. Once the forepaw digits were seen to be wrapped around the bar, mice were pulled laterally, parallel with the benchtop, until grip on the bar was broken. Five to fifteen pulls (measured in pound-force, LBF) were collected, depending on the consistency of performance, in groups of roughly five pulls. Pulls were averaged, then analyzed.

#### Rotarod

Groups of up to five mice were tested in the Accelerating Rota-Rod for Mice (Jones & Roberts model 7650; Ugo Basile), each assigned a lane per session. Mice were trained before each testing session at roughly 4 rotations/minute (rpm) with no acceleration for three, 1-minute intervals (each separated by 2 minutes of rest in their home cages). Mice were then tested by an accelerating protocol, from ∼4 rpm to ∼40 rpm, over 5 minutes. The latency was measured for each mouse to either fall off the rod or passively cling to/rotate with the rod for two consecutive rotations. Each test consisted of five trials, each separated by 2 minutes of rest in their home cages. The three trials with the longest latencies were then averaged and analyzed. The entire apparatus was thoroughly cleaned with 70% ethanol between groups of mice.

#### Gait Analysis

Gait analysis was completed using the DigiGait (Mouse Specifics Inc.) treadmill and software. Mice were placed in the walking chamber and acclimated to the task at a treadmill speed of 15 cm/s. The treadmill was then paused, the speed increased to 20 cm/s, then turned back on for testing. After the mouse was freely moving and fluidly walking, 6–12 seconds of video was recorded. Any excrement or urine on the treadmill belt was wiped off with 70% ethanol between mice.

Video preparation followed Digigait software workflow, including pre-processing, processing, and post-processing. During pre-processing, paw detection was optimized using contrast functions to digitally “paint” the paws. During processing, paw positions were tracked frame by frame, generating dynamic gait signals representing the paw contact area of each limb over time. In post-processing, each gait signal was reviewed alongside video playback to confirm traces accurately reflected limb movement. Periods during which mice remained stationary, sat on the treadmill belt or back bumper rail, grabbed the front bumper rail, or reared were excluded from analysis. Finalized traces, were analyzed using the Digigait software, exported, and compiled for statistical analysis.

Left and right hindlimb data were averaged to produce a single hindlimb score per metric, per mouse. The variables chosen for gait analysis were Stride, Swing, Stance, Stance/Swing, Propel, and Paw Drag. Stride, the duration of one complete stride for a particular paw, is made up of two phases: Swing (the time during which a paw is not in contact with the ground) and Stance (the time during which the paw is in contact with the ground). Stance/Swing is the ratio of these phases. Propel is the time the paw spends pushing off of the ground, ie the time from the moment of maximum paw contact to the moment it is completely lifted off the ground. Paw drag, a metric describing propulsion, is the area under the curve from the moment of maximal stance to the moment the paw lifts off the ground^58^.

### Cognitive and Behavioral Measures

#### Open Field Assay

For cognitive assessments, up to four mice were tested at a time, each in an acrylic arena, measuring 40cm x 40cm x 30cm. Each arena was wrapped in aluminum foil to prevent mice from seeing neighboring arenas. During the Open Field Assay, mice were allowed to behave in their assigned arena freely for 10 minutes. Video was recorded for all four arenas by a camera mounted on the ceiling above the arenas and analyzed using Any-maze Video Tracking System (version 7.6) Software.

### Pathology

#### Harvesting

Mice were euthanized with carbon dioxide in accordance with IACUC regulations and perfused transcardially with ice-cold phosphate buffered saline (PBS) to clear residual blood. For *C9orf72* repeat expression analysis across the CNS and peripheral organs, tissues (cerebral cortex, cerebellum, spinal cord, and spleen) were collected immediately following PBS perfusion and flash frozen in liquid nitrogen. For *C9orf72* repeat expression analysis across the cervical spinal cord and all immunohistochemistry assessments, animals underwent PBS perfusion followed by trasncardiac perfusion with 4% paraformaldehyde (PFA) in PBS for tissue fixation. Both the spinal cord and brain were dissected, post-fixed overnight in 4% PFA at 4°C, paraffinized using the Tissue-Tek VIP Tissue Processor system (Miles/Sakura), then embedded in paraffin. For GFP fluorescent analyses, animals were similarly perfused with PBS followed by 4% PFA. Spinal cord and brain were dissected and post-fixed overnight in 4% PFA. After post-fixation, tissue was transferred to a 30% sucrose cryoprotectant solution and incubated at 4°C. Tissue was then embedded and frozen in blocks of Tissue-Tek OCT Compound (Sakura).

#### Expression (C9-NRE and GFP)

For qPCR expression analysis across the CNS and organs, Trizol extraction was performed from fresh frozen tissue of PBS perfused animals. For cervical spinal cord expression analysis, RNA was isolated from upper cervical sections of PBS and PFA perfused animals. Tissue was paraffin embedded, microtome-cut, and processed with the FFPE PureLink RNA Mini Kit (ThermoFisher). All mRNA, regardless of extraction method, was reverse transcribed using the QuantiTect Reverse Transcription Kit (Qiagen) and was then prepared for qPCR using a SYBR Green qPCR master mix (ThermoFisher). qPCR runs and analysis were performed on a QuantStudio 5 Real-Time PCR System (ThermoFisher). Samples were measured in technical triplicate from each animal for both *C9orf72* and GAPDH gene levels. Data are represented as fold change comparisons between *C9orf72* repeat expression and GAPDH, per animal. Primer sets were obtained from Integrated DNA Technologies (Coralville, IA). Primer pairs for the *C9orf72* repeat were previously described by Zhu et al (2020):

C9 Repeat Forward: AGCTTAGTACTCGCTGAGGGTG

C9 Repeat Reverse: GACTCCTGAGTTCCAGAGCTTG

Mouse GAPDH Forward: AACAGCAACTCCCACTCTTC

Mouse GAPDH Reverse: CCTGTTGCTGTAAGCCGTATT

#### Immunohistochemistry (IHC) Sectioning

Paraffin embedded tissue was cut on a microtome at a thickness of 10μm. For staining, starting at the cervical-thoracic border of the spinal cord, two sections were collected per slide. 255μm deeper into the block/further rostral in the cervical region, two more sections were collected, yielding four sections from two regions per slide. This process was repeated once more per cervical spinal cord to yield a total of eight sections, across four regions, spanning roughly 1.5–2.0mm, depending on the number of slides needed per mouse. Given the anatomical organization of the spinal cord, this method of sectioning allowed for pathological analysis across a functional range of the cervical spinal cord, capturing pathology roughly associated with C6–T1, including the motor regions associated with forelimb control.

#### Immunofluorescent (IF) Staining

Frozen, OCT-embedded tissue was cryo-sectioned at a thickness of 20μm, placed on slides, and dried at room temperature overnight. For staining, slides were washed in PBS, permeabilized with a Triton solution, then treated with a BSA protein block solution in a hydrated chamber. Primary antibodies were then applied to slides, incubating overnight at 4°C. After rinsing the following day, slides were incubated in secondary antibodies at [1:1000], at room temperature, then slides were rinsed again. Hoechst was added to the penultimate rinse then slides were coverslipped using ProLong Diamond Antifade Mountant (Invitrogen P36961). Antibodies and concentrations were as follows: NeuN (Cell Signaling D4G4O) [1:1000]; GFP (Abcam ab13970) [1:1000]; Hoechst 33342 (Invitrogen H1399) [1.67 ug/mL].

#### IHC Staining

Slides undergoing IHC staining were first deparaffinized using the Gemini AS system (Thermo Scientific), moving from xylene to decreasing concentrations of ethanol solutions to water. Slides were exposed to antigen retrieval in a heated, high pH solution (Invitrogen 00-4956-58), then rinsed and treated with a BSA protein block solution in a hydrated chamber. Primary antibodies, diluted in the same protein block solution, were then applied to slides, incubating overnight at 4°C. On the second staining day, after rinsing, the Vectastain Elite ABC Kit (Rabbit IgG; PK-6101) was used, moving slides through the system from Biotinylated Secondary antibody, rinse, Elite ABC Solution, and rinse. Slides were developed using peroxidase detection solution from the DAB Substrate Kit (Vector Labs SK-4100), then plunged into water to stop the reaction. Hematoxylin counterstain (Sigma-Aldrich GHS332) followed by an ethanol/hydrochloric acid differentiation was applied. Slides were then dehydrated, again using the Gemini AS system (Thermo Scientific), moving from water to increasing concentrations of ethanol solutions to xylene, then coverslipped with ClearVue Mountant XYL (Thermo Scientific 4212).

In addition to the above “base” staining protocol, after deparaffinization, poly-GP, poly-GR, and pTDP-43 slides were incubated in an additional blocking solution of 5% hydrogen peroxide in methanol, then rinsed. While PBS was used for all other stains, pTDP-43- and poly-GA-stained slides used TBS. NeuN and GFAP were co-stained, with NeuN staining following the base protocol over the first two days of staining, through DAB development. After development, slides underwent BSA protein block again, and were then treated with the second/GFAP primary antibody, incubating overnight at 4°C. On the third day of staining, slides were rinsed, treated with the Vectastain Elite ABC Kit (Rabbit IgG; PK-6101) but with substituted Biotinylated Rat IgG (BA-9401-.5), and then developed using the VIP Substrate Kit (Vector Labs SK-4600).

After development, the NeuN-GFAP slides progressed through counterstaining, dehydration, and coverslipping (differentiation was incompatible with the VIP stain, and therefore skipped). Finally, for poly-GA staining, antigen retrieval was performed in IHC-Tek epitope retrieval solution (IHC World IW-1100-1L). Endogenous peroxidase was blocked by incubating with BLOXALL (Vector Labs SP-6000-100). Tissues underwent blocking in Protein block (DAKO X090930-2), then were incubated with primary antibody overnight. The next day, DAKO Envision +HRP polymer kit (K4003) was used, and the reaction was visualized using ImmPACT VIP Substrate Kit (Vector Lab SK-4605). Sections were then mounted with Aqua-Poly-Mount (Polysciences 18606-20). Primary antibodies and concentrations were as follows: GFAP (Invitrogen 13-0300) [1:5,000]; GP Repeats (Proteintech 24494-1-AP) [1:25,000]; GR Repeats (Proteintech 23978-1-AP) [1:7,000]; GA Repeats (gift of Dr. Leonard Petrucelli^59^) [1:50,000]; Iba1 (Wako 019-19741) [1:10,000]; NeuN (Cell Signaling D4G4O) [1:3,000]; pTDP-43 Ser409/410 (Proteintech 22309-1-AP) [1:60,000].

#### Imaging and Image Processing

Imaging of IF staining was performed on a Nikon A1^+^ confocal microscope and a Nikon Eclipse Ti microscope, using NIS-Elements software. The cortex was imaged at 10× (Nikon CFI Plan Fluor 10× objective MRH10105), the cerebellum and spinal cord cross-sections were imaged at 20× (Nikon CFI Plan Apo λ 20× objective MRD00205), and the spinal ventral horn regions, primary motor area, and cerebellar Purkinje cells were imaged at 60× (Nikon CFI Plan Apo Lambda 60× Oil Immersion objective MRD01605; Figure 1B, Figure S1).

**Figure 1:**
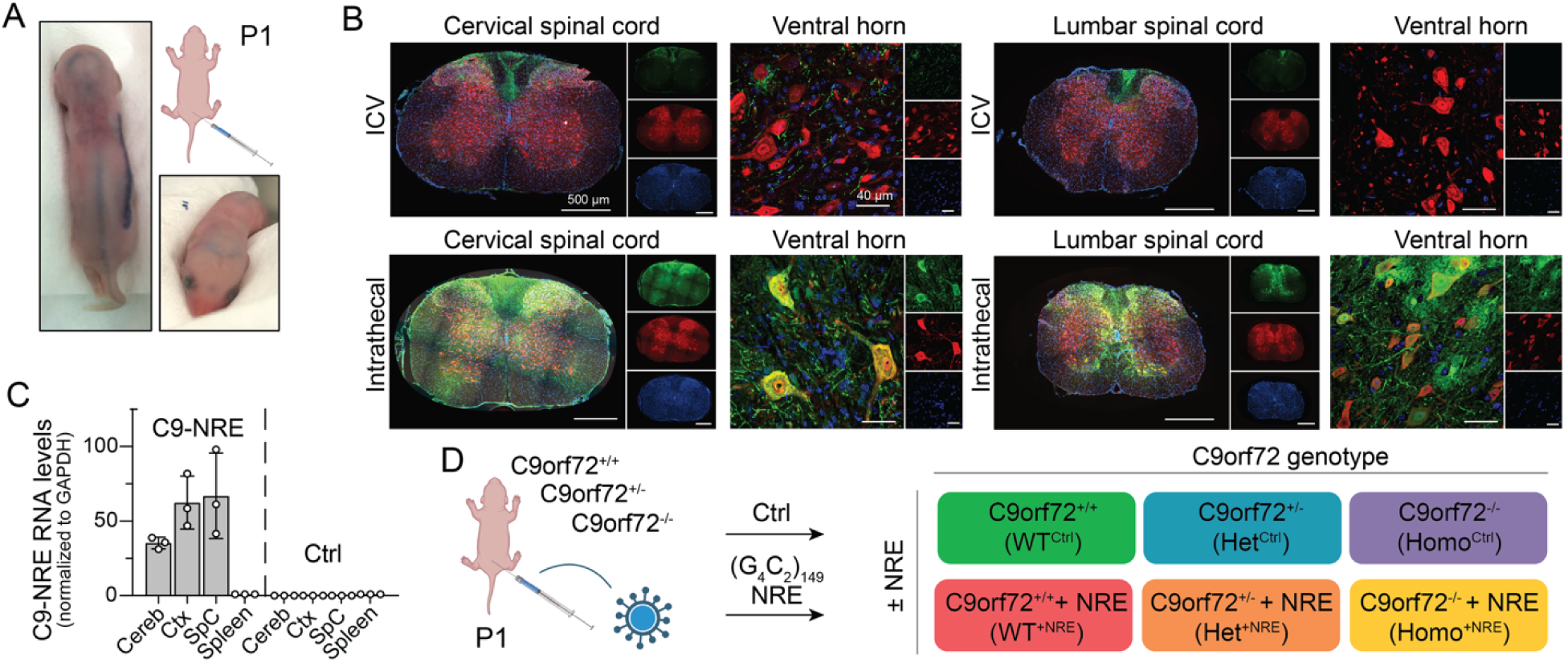
Intrathecal AAV delivery achieves widespread (G_4_C_2_)_149_ repeat expression throughout the CNS, including robust spinal cord engagement. (A) Intrathecal injection of a P1 pup shows injectant rapidly distributes throughout the CNS. (B) Intrathecal injections of AAV2/9 GFP reporter show robust expression throughout the spinal cord at four weeks post-injection. Green: GFP; Red: NeuN; Blue: Hoechst. (C) RT-qPCR detects (G_4_C_2_)_149_ repeat expression in tissues consistent with the GFP reporter expression patterns in (B), and is not detected in the periphery or in Controls at four weeks post-injection. Expression is normalized to GAPDH. Bars represent mean ± SD; *n* = 3. (D) Intrathecal injections of AAV2/9 (G_4_C_2_)_149_ into *C9orf72* knockout background mice yields cohorts either with or without C9-NRE expression, and either full (WT), partial (Het), or complete knockout (Homo) of *C9orf72* expression to appreciate both gain- and loss-of-function mechanisms, with pathophysiologically relevant cell and regional C9-NRE AAV engagement.

Fluorescent images of the brain were processed using Fiji, including thresholding for background fluorescence compared to controls and for large, stitched images, background subtraction using a 50-pixel rolling ball radius. The green channel pictured displays the GFP reporter signal. Fluorescent images of the spinal cord were also processed using Fiji, again thresholding for background fluorescence compared to controls. The green channel pictured displays the GFP reporter combined with the GFP antibody signal.

For most IHC, whole-slide, brightfield images were acquired by the Jefferson Translational Pathology Shared Resource Core using a 3DHistech P1000 digital scanner at 20× magnification. IHC images were then pixel thresholded against background stain using QuPath (version 0.6.0), then compiled for analysis. For poly-GA IHC, whole-slide, brightfield images were acquired as 20× magnification tiles using Zeiss Axio Imager and analyzed using ZEN (version 3.4). To improve visualization and highlight regions of interest, 20× images were digitally enlarged for figure presentation by 10× (Figure 7A), 5× (Figure 7D), and 13× (Figure 7G and Figure S6).

For the quantification of all IHC except pTDP-43, the ventral horns were manually traced as the region of interest (ROI). One ROI per each of the four cervical regions sectioned was subjected to analysis to ensure a unique population of cells was analyzed per region. Pixel-level segmentation was performed using predefined optical density and color feature thresholds. Threshold values were determined using representative images and then applied unchanged in batch to all images of the same target to identify positively stained regions and exclude background. Similar to previous studies^29,45,46^, data were normalized by the area of the associated ventral horn and then averaged across the four cervical regions to produce a normalized area of positive stain per animal. Due to the sparse nature of pTDP-43 pathology, the entire grey matter was traced as the ROI and considered for analysis. Additionally, both sections per region were analyzed, resulting in data from eight sections’ grey matter (rather than four sections’ ventral horns for the other targets) contributing to each animal’s assessment. Rare instances where sections displayed evidence of technical processing issues or were damaged during the staining process were not included in analyses.

### Data and Statistical Analyses

#### CMAP Signal Preprocessing and Machine Learning-Based Feature Prediction

Raw CMAP traces were acquired from text files output by the Neurosoft Neuro-MEPomega software and parsed using custom R scripts (R version 4.3.2). Baseline correction was performed per trace using the mean signal within a window from 15 to 20 ms. To evaluate electrophysiological parameters, we implemented a machine learning pipeline using the ranger package for random forest regression. First, engineered signal features were derived from each trace, including Amplitude (peak-to-peak voltage) and Absolute Area Under the Curve (AUC) when ≥50 labeled observations were available. These features served as predictors for training individual random forest models for each physiological target variable. All models were trained starting with 500 trees. Predictions from these models were used to generate a predicted AUC for each trace. All analyses were performed in R using standard packages.

#### Survival Analysis

Mice not harvested for 6- or 9-month IHC analyses were tracked longitudinally for survival. Survival curves were generated using the Kaplan–Meier method and compared using log-rank (Mantel–Cox) and Gehan–Breslow–Wilcoxon tests in Prism (v10.6.1). Animals harvested for 12-month IHC and expression analyses were censored at the time of tissue collection. A log-rank test for trend was also applied across genotypes and treatment groups (WT^Ctrl^, Het^Ctrl^, Homo^Ctrl^, WT^+NRE^, Het^+NRE^, Homo^+NRE^).

#### Mixed Model Analysis

Motor Performance and Behavioral Measures data were initially assessed at the latest time point (12- and 10-months, respectively) using Prism (version 10.6.1) Mixed Model analyses across the fixed effects and interactions of Genotype (WT, Het, Homo), Treatment (Ctrl, +NRE), and Sex (male, female) to direct further analysis. Variables that failed the Shapiro–Wilk test for normality (*p* value < 0.05) were re-analyzed using IBM SPSS Statistics (version 31.0.1.0) Generalized Linear Mixed Models, applying both Linear and Gamma Target Distributions. In these cases, the model with the lowest Akaike Information Criterion, indicating the better fitting model, was chosen. Both Rotarod and Open Field-Total time mobile were only analyzed targeting a Gamma Distribution, due to the maximum being dictated/cut off as the total time of each experiment. Open Field-Immobile episodes was assessed according to a Poisson Distribution, as is appropriate for count-based data. For variables where no significant differences were found across Sex, further analyses were carried out combining data from males and females. For variables were there was a significant difference found across Sex, subsequent analyses were carried out for males and females separately. 12- and 10-month statistics can be found in Table S1.

Following this initial assessment, Gait and Open Field data was assessed using Prism Mixed Model analyses across the fixed effects and interactions of Genotype and Treatment (Table S1 (Single time point)). Again, any variable that failed the Shapiro–Wilk test for normality was re-analyzed using IBM SPSS Statistics Generalized Linear Mixed Models, applying both Linear and Gamma Target Distributions, choosing the model with the lowest Akaike Information Criterion. Again, Open Field-Total time mobile was targeted to a Gamma Distribution and Open Field-Immobile episodes was targeted to a Poisson Distribution due to the nature of each experiment.

After the initial assessment, Electrophysiology, Grip Strength, Rotarod, and Body Mass data were assessed using IBM SPSS Statistics Generalized Linear Mixed Models, applying both Linear and Gamma Target Distributions (choosing the model with the lowest Akaike Information Criterion; except in the case of Rotarod-again, assessed targeting only a Gamma Distribution) across the Fixed Effects and interactions of Genotype, Treatment, and Time (Table S1 (Longitudinal)).

IHC data was similarly analyzed using IBM SPSS Statistics Generalized Linear Mixed Model, applying a Gamma Target Distribution. Fixed Effects and interactions (when appropriate) of Genotype and Treatment were assessed for poly-GP, poly-GR, NeuN, GFAP, and Iba1. Fixed Effects and interactions of Treatment and Time were considered for poly-GP, poly-GR, and Iba1 across time (Table S1).

For all IBM SPSS statistical analyses, the Random Effect of Ear_Tag (aka subject) was included to improve model fit. If the addition of the Random Effect interfered with the convergence of the model, it was excluded.

#### Correlation Analyses

To examine relationships across motor performance, behavioral measures, and pathological domains, correlation analyses were performed using a curated set of experiment–variable combinations selected a priori. For each mouse, measurements within a given experiment and variable were averaged, when applicable, to yield a single value per feature, and features were then assembled into a per-animal matrix. Analyses used the last acquired time point.

Pairwise associations between numeric variables were assessed using Pearson’s correlation with pairwise complete observations. Two-sided *p* values were computed for each feature pair when at least three complete observations were available. Additionally, to account for multiple comparisons, *p* values were adjusted using the Benjamini–Hochberg false discovery rate (FDR) procedure.

For follow-up visualization of significant phenotype–phenotype relationships, feature pairs with FDR-adjusted *p* < 0.05 were selected from the dot-plot results. A simple least-squares linear model was fit to all mice with complete data for that pair, and R² and *p* values were extracted; *p* values across plotted pairs were additionally FDR-adjusted using the Benjamini–Hochberg procedure.

## RESULTS

### Neonatal intrathecal (G_4_C_2_)_149_ AAV injections express expanded repeats throughout the brain and spinal cord

To achieve C9-NRE expression throughout the central nervous system, including spinal cord regions implicated in ALS/FTD, we leveraged intrathecal delivery approaches shown to support both brain and spinal cord engagement. Previous studies have achieved C9-NRE expression throughout the brain using AAV2/9 delivery via ICV injection into neonatal pups; however, this approach has shown limited spinal cord expression. In contrast, intrathecal delivery of other AAV2/9 systems have reported both brain and spinal cord expression, including engagement of neurons, astrocytes, and motor neurons in the spinal cord^50,52^. Therefore, to enhance spinal cord expression while preserving engagement of brain regions targeted by prior C9-NRE ICV studies^29,45^, we performed intrathecal injections of the equivalent AAV2/9-GFP reporter on P1 pups. Immediately following injections, dye from the injectant was clearly visible throughout the spinal cord, as well as in the cerebellum, along the cranial sutures, and in the forebrain/olfactory region (Figure 1A). Four weeks post-intrathecal injection, robust GFP expression was observed throughout the spinal cord, extending from the lumbar to cervical levels. Notably, GFP signal was present in NeuN cells located throughout the cord, including large, pyramidal neurons in the ventral horn. In contrast, comparable ICV injections showed limited cervical expression and qualitatively undetectable GFP expression in the lumbar region (Figure 1B). In addition to spinal cord expression, intrathecal injections produced strong GFP signal in multiple brain regions, including cortex, hippocampus, and cerebellum, with labeling observed in NeuN cells within the primary motor region and in cerebellar Purkinje cells (Figure S1A–B). Overall, GFP spinal cord expression is consistent with previously reported AAV2/9 intrathecal injection studies^50,52^ and the distribution of brain expression is qualitatively comparable to that described in prior studies using ICV delivery of C9-NRE AAV vectors^29,45^.

Based on these GFP expression patterns, we next examined C9-NRE expression following intrathecal injection. P1 pups were intrathecally injected with a previously characterized^44^ AAV2/9 vector encoding a (G_4_C_2_)_149_ C9-NRE or dye only (Control). C9-NRE expression was quantified by qPCR in cortex, cerebellum, spinal cord, and spleen, normalized to GAPDH. Four weeks post-injection, repeat expression was readily detected throughout the CNS, while no expression was detected in the spleens of repeat-injected animals or in any tissues from Control mice (Figure 1C).

The AAV dose and control conditions used throughout this study were guided by prior intrathecal and neurodegenerative AAV studies^48–54^, which demonstrated that comparable dosing^54^ of an AAV-GFP Control does not induce motor deficits, neurodegeneration, or transcriptome alterations, and that there were no differences between sham and non-pathogenic (G_4_C_2_)_2_ AAV-injected animals when using a similar vector^45^. Together, these findings support attribution of the phenotypes described here to pathogenic C9-NRE expression rather than AAV delivery.

Capitalizing on this delivery strategy, P1 pups from a well-characterized *C9orf72* knockout line^40^ were subsequently intrathecally injected to generate six experimental cohorts defined by C9-NRE expression (+NRE or Ctrl) and endogenous *C9orf72* genotype (WT, Het, or Homo; Figure 1D). This design enabled systematic evaluation of the independent and combined effects of C9-NRE expression and *C9orf72* genotype across longitudinal motor, behavioral, and pathological measures relevant to ALS/FTD.

### C9-NRE expression drives progressive deficits in muscle strength and motor unit function

Given the central role of motor dysfunction in ALS/FTD and the variable effects on muscle strength reported in prior C9-NRE mouse studies^29,44,45^, we examined muscle strength as a functional outcome of CNS-wide C9-NRE expression, including the spinal cord, across *C9orf72* genotypes. Forelimb grip strength was assessed longitudinally, with initial analyses performed at a later time point to account for the age-dependent emergence and progressive nature of ALS/FTD phenotypes, as well as the absence of significant survival differences despite a log-rank trend (Figure 2A; Figure S2A). At 12 months of age, grip strength was significantly influenced by genotype (*p* = 0.003), C9-NRE expression (*p* < 0.001), and sex (*p* < 0.001) (Figure 2B; Table S1). Pairwise contrasts showed reduced grip strength in C9-NRE mice compared to Controls (*p* < 0.001), and that Homo mice were weaker than both WT and Het mice (*p*_WT vs Homo_ = 0.001; *p*_Het vs Homo_ = 0.010). Male mice displayed higher grip strength than females, consistent with known sex-dependent differences in body mass. Accordingly, body mass was examined prior to further analysis (Figure S2B–C). As expected, body mass was strongly dependent on sex (*p* < 0.001), but was also significantly reduced by C9-NRE expression (*p* < 0.001). Because repeat-associated reductions in body mass (i.e., potential muscle atrophy) could contribute to decreased grip strength, subsequent grip strength analyses were stratified by sex rather than normalized to body mass (Figure S2D).

**Figure 2:**
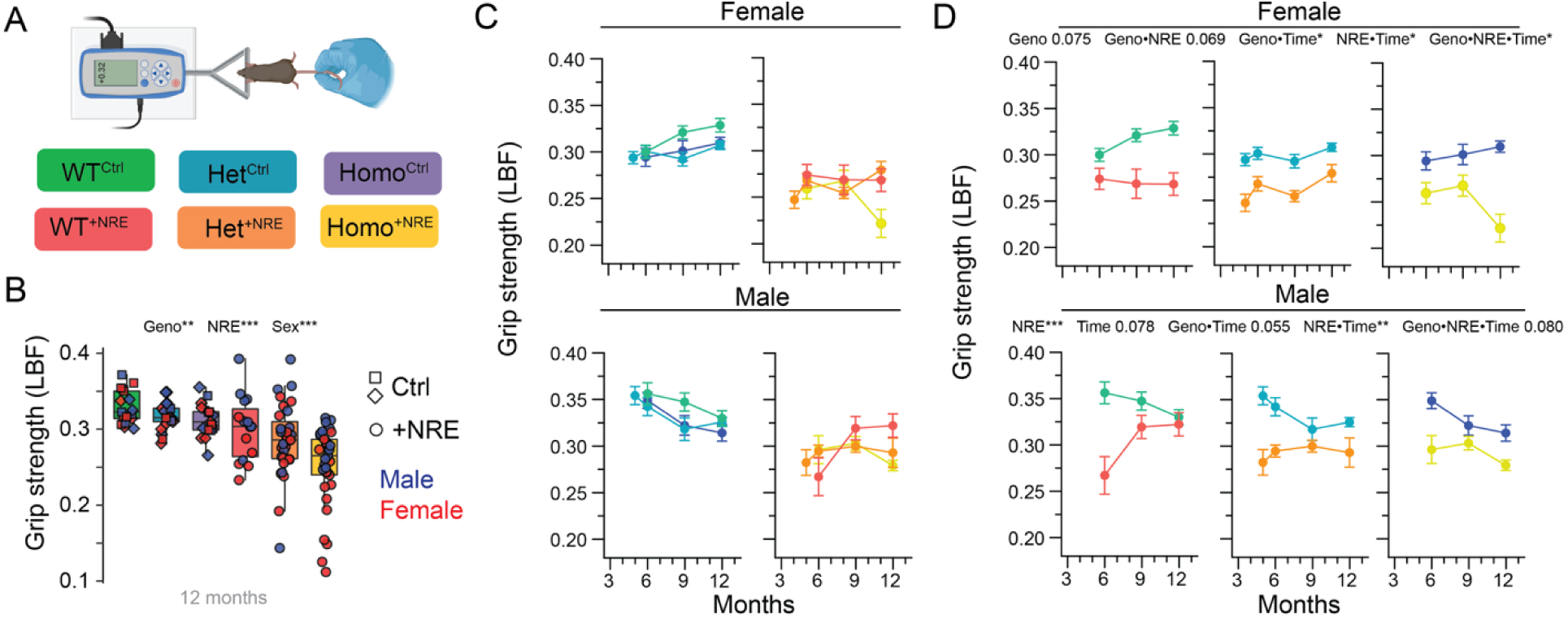
C9-NRE (G_4_C_2_)_149_ expression reduces grip strength. (A) Schematic of grip strength testing with triangle pull. (B) At 12 months of age, grip strength is reduced in C9-NRE mice compared to Controls and in Homo mice compared to both WT and Het mice. Also, males exhibit higher grip strength than females. (C–D) Longitudinal grip strength data grouped first by C9-NRE (C), then by genotype (D), show reduced grip in C9-NRE mice, regardless of genotype, with additional deficits observed in Homo^+NRE^ mice at later time points. Box plots are Tukey box plots showing median, Q1 and Q3, and whiskers at 1.5× interquartile range. Line graphs show mean ± SEM. *n* = 5–25 per group, defined by ± C9-NRE expression, *C9orf72* genotype, sex, and time (e.g., Female Het^+NRE^ mice at 6 months).

Extending the analysis across the entire time course to capture disease progression revealed C9-NRE expression as the predominant driver of grip strength differences. Both female and male mice showed significant differences in C9-NRE × time interactions (*p*_Female_NRE×Time_ = 0.022; *p*_Male_NRE×Time_ = 0.002), indicating grip strength trajectories differ significantly between Control and C9-NRE groups (Figure 2C–D). Pairwise contrasts corroborated this pattern, with C9-NRE mice exhibiting lower grip strength than Controls both overall and within genotype groups (Figure 2C–D). The interaction of genotype × C9-NRE did not reach significance in either sex (*p*_Female_Geno×NRE_ = 0.069; *p*_Male_Geno×NRE_ = 0.100), consistent with overlapping grip strength ranges within the Control and C9-NRE groups (Figure 2C). Genotype × time and genotype × C9-NRE × time interactions were significant or approached significance in both sexes, indicating that genotype-dependent divergence emerges at later time points. Further examination of longitudinal trajectories, together with the 12-month analysis, suggest that these effects are driven by reduced grip strength in Homo^+NRE^ mice at 12 months (Figure 2D). Collectively, these data indicate that C9-NRE expression is the driving factor underlying progressive grip strength decline, and that loss of *C9orf72* function contributes additional effects that become more apparent with aging.

To determine whether motor dysfunction was evident at the level of motor unit output, CMAPs from the footpad muscles were analyzed following stimulations at the sciatic notch (hip region) and the ankle (Figure 3A). Both amplitude (peak to peak) and area (absolute area under the curve) were assessed (Figure 3B). At 12 months of age, CMAP amplitude was significantly influenced by C9-NRE expression and sex at both stimulation sites (all *p* < 0.001), with repeat-expressing mice exhibiting lower amplitudes than Controls and males exhibiting higher amplitudes than females (Figure 3C). No significant main effects of genotype or genotype-related interactions were detected at this time point.

**Figure 3:**
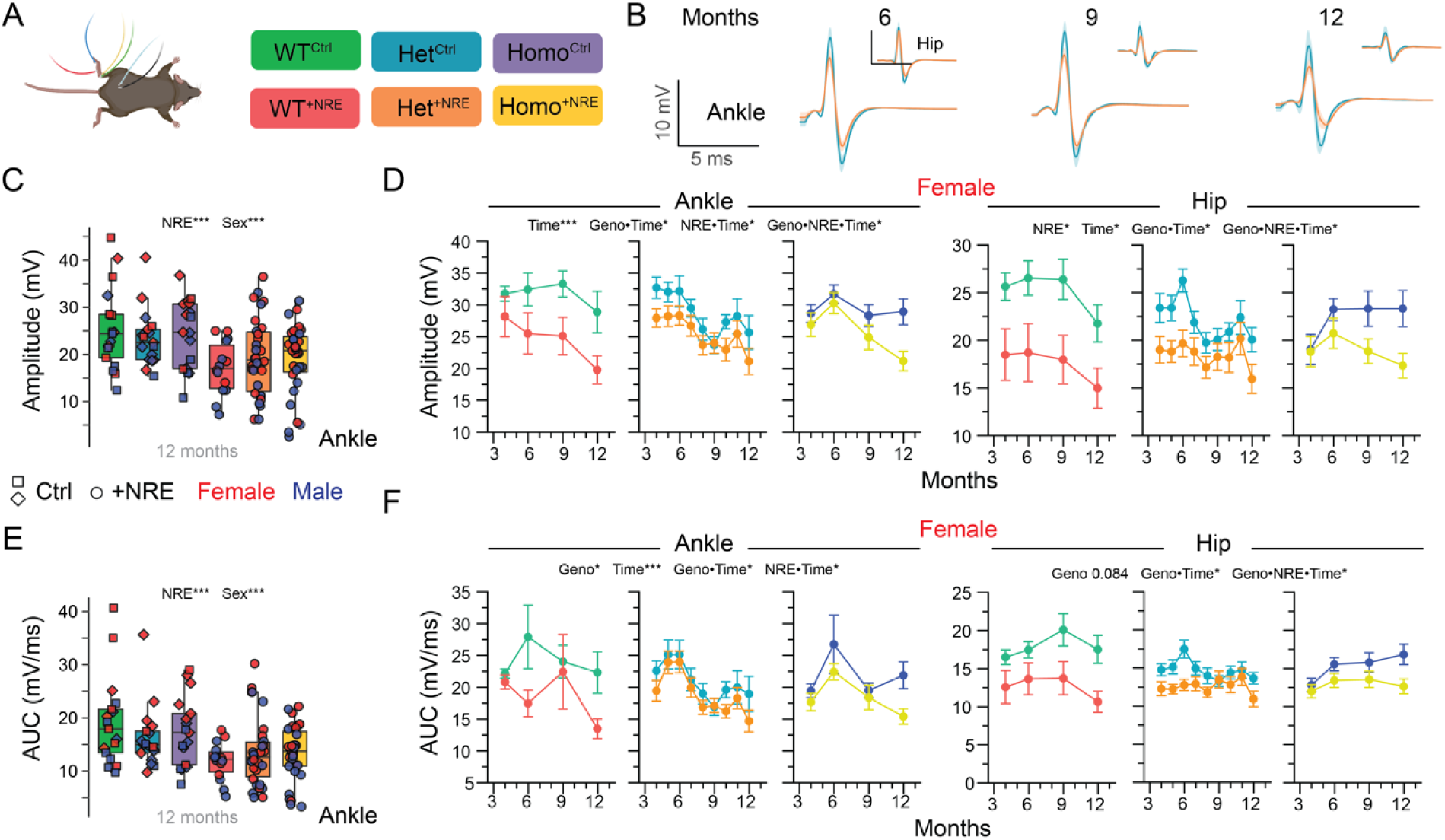
Electrophysiological deficits in C9-NRE (G_4_C_2_)_149_-expressing mice reflect mild ALS-like phenotype. (A) Schematic of electrophysiology set up with stimulation sites at the sciatic notch and ankle and recording electrodes at the footpad muscles. (B) Average traces for Het^+NRE^ (orange) and Het^Ctrl^ (teal) groups at 6-, 9-, and 12-months with SEM (corresponding color shadows), illustrating progressive reductions in CMAP amplitude and area. (C) At 12 months, C9-NRE mice have lower amplitudes than Control mice. Sex is also a significant factor dictating CMAP amplitude. (D) Longitudinal analysis grouped by genotype shows reduced CMAP amplitudes in C9-NRE mice compared to Controls. (E) At 12 months, CMAP areas differ according to C9-NRE and sex. (F) Repeat-containing mice also show decreases in CMAP area compared to Controls. Together, decreases in both amplitude (D) and area (F) suggest potential axonal loss and/or muscle atrophy. Box plots are Tukey box plots and line graphs show mean ± SEM. *n* = 5–24 per group.

Based on these findings, CMAP amplitude was next examined longitudinally with sex-stratified analyses to assess the contributions of C9-NRE expression, genotype, and aging. Across stimulation sites and sexes, time had a significant effect on CMAP amplitude, consistent with age-related deficits in motor unit function (Figure 3B, 3D; Figure S3A). In multiple site- and sex-specific comparisons, significant C9-NRE × time interactions were observed (*p*FemaleAnkle_NRE×Time = 0.023; *p*MaleAnkle_NRE×Time < 0.001; *p*MaleHip_NRE×Time = 0.011), indicating that CMAP amplitude declines more steeply over time in repeat-expressing mice compared with Controls. Genotype-related effects were limited overall, as reflected by substantial overlap among C9-NRE groups and among Control groups across genotypes. However, in female mice, genotype interactions emerged at later time points, including significant genotype × C9-NRE × time interactions for both ankle and hip stimulation (*p* = 0.032 and *p* = 0.047, respectively). Visual inspection of longitudinal trajectories indicates that these effects are driven by divergence between Homo^+NRE^ and Homo^Ctrl^ mice at later time points (Figure 3D). Together, these findings indicate that C9-NRE expression drives progressive CMAP amplitude decline, with genotype-dependent effects becoming detectable over time in a sex-specific manner.

Similar patterns were observed when CMAP area was examined. At 12 months of age, CMAP area was significantly influenced by C9-NRE expression and sex (all *p* < 0.001), while no genotype-related differences were detected (Figure 3E). Longitudinal analysis revealed greater variability in the effects of C9-NRE across stimulation sites and sexes compared with CMAP amplitude (Figure 3B, 3F; Figure S3B). Time remained a significant factor, indicating that CMAP area decreases with aging, and several site- and sex-specific comparisons showed significant or near-significant differences driven by C9-NRE, reflected as differences in the range or trajectory of CMAP area between repeat-expressing mice and Controls. As with CMAP amplitude, genotype-related effects were most evident in female mice, including significant genotype × C9-NRE × time interactions following hip stimulation, but pairwise contrasts consistently highlighted reduced CMAP area in C9-NRE mice compared with Controls, including within genotype groups. Overall, CMAP amplitude and area data show a concordant, progressive decline consistent with ongoing motor unit dysfunction driven by C9-NRE expression, with genotype-dependent effects emerging over time in a sex-specific manner, potentially exacerbating motor impairment.

In aggregate, forelimb grip strength and sciatic-footpad electrophysiology demonstrate decreased muscle strength in C9-NRE mice, supporting repeat expansion gain-of-toxicity mechanisms playing a role in motor performance. They also indicate that *C9orf72* genotype, particularly homozygous loss of function, might play a delayed or synergistic role in the deterioration of muscle function.

### Decreased rotarod performance is primarily *C9orf72* genotype-dependent

Given mixed reports of motor phenotypes in prior C9-NRE ICV models, where gain-of-function approaches yielded variable outcomes^43,45^ but a combinatorial model showed impaired rotarod performance^29^, we assessed motor function using rotarod testing, a composite measure that predominantly reflects motor coordination. Performance was quantified as latency to fall or passive rotation with the rod (Figure 4A). At 12 months of age, genotype had a significant effect on rotarod performance (*p* < 0.001), whereas C9-NRE expression and sex were not significant contributors (Figure 4B). Pairwise contrasts revealed that while Het mice exhibit reduced latencies compared to WT mice, these groups were not statistically different. Homo mice, however, displayed significantly reduced rotarod performances relative to both WT and Het mice (both *p* < 0.001; Figure 4B).

**Figure 4:**
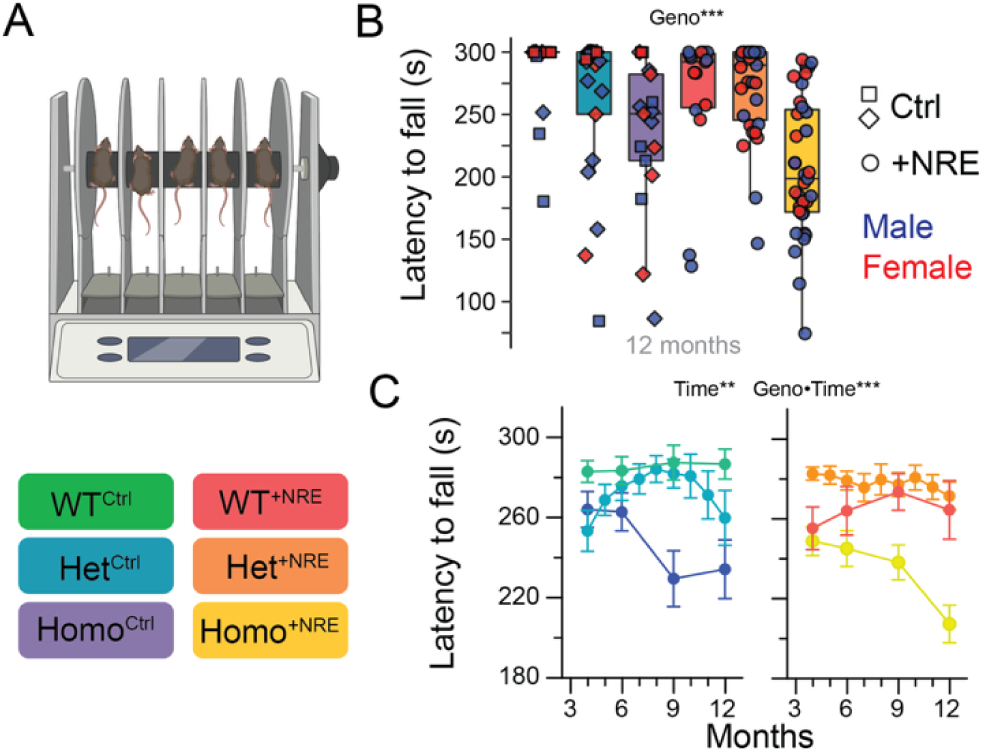
*C9orf72* genotype predominantly drives rotarod deficits. (A) Schematic of accelerating rotarod. (B) Latency to fall at 12 months of age is genotype-dependent. (C) Rotarod data, grouped here by C9-NRE expression, show reduced performance based on genotype regardless of repeat expansion. Homo mice produce shorter latencies to fall compared to their Het and WT counterparts, regardless of the presence of C9-NRE. Box plots are Tukey box plots and line graphs show mean ± SEM. *n* = 6–25 per group.

Extending the analysis across all time points to evaluate disease progression revealed similar, genotype-dependent effects on motor coordination. While time, and therefore general aging, had a significant effect on rotarod performance (*p* = 0.006), C9-NRE expression and all C9-NRE-related interactions did not significantly influence rotarod performance, indicating that repeat expression did not measurably alter motor coordination (Figure 4C; Figure S3C). Interestingly, a significant genotype × time interaction was observed (*p* < 0.001), meaning that the trajectories of rotarod performance differed among genotypes (Figure 4C). Both WT and Het mice exhibited relatively stable performances from 4 to 12 months, with no significant differences detected between these groups. However, Homo mice showed pronounced instability, sharply declining in rotarod performance after 6 months of age, and pairwise contrasts confirmed significantly reduced latencies compared with WT and Het mice (*p* = 0.001 and *p* < 0.001, respectively; Figure 4C; Figure S3C), indicating Homo mice are largely responsible for the genotype × time differences. Notably, pairwise contrasts within the Homo mice revealed differences between Control and C9-NRE mice approaching significance (*p* = 0.072; Figure 4C). Combined with a similar trend at 12 months (*p* = 0.093; Figure 4B), these data indicate that C9-NRE effects on rotarod performance may emerge or become more pronounced at later time points, potentially interacting with genotype-dependent motor coordination deficits.

Together, these data demonstrate that the progressive decline in motor coordination is predominantly driven by *C9orf72* genotype, particularly by homozygous loss-of-function.

### Gait analysis identifies subtle signatures of motor dysfunction

As a complementary assessment of motor coordination and function, we also examined whether this model exhibits disease-relevant alterations in gait (Figure 5A). Gait analysis has been widely used to detect subtle and progressive motor abnormalities in patients and preclinical ALS models^60–62^, and was assessed here using the Digigait system. To unbiasedly reduce the dimensionality of the dataset, we applied principal component analysis (PCA) to fore- and hindlimb gait measures at 12 months of age (Figure S4A). This analysis identified a subset of parameters contributing most strongly to variance across experimental groups and focused subsequent analyses on hindlimb measures. Among these parameters: Stride, Stance, Propel, and Paw Drag were all significantly influenced by genotype; Stance/Swing was significantly affected by C9-NRE expression; and Swing was influenced by both genotype and repeat expansion (Figure 5B–D; Figure S4B–D). Only Propel showed differences across the main effect of sex, possibly due to the body mass differences between sexes, therefore, the other gait parameters were pooled across sex in further analyses. Together, genotype-driven differences in Stride (*p* = 0.011), Swing (*p* = 0.027), and Stance (*p* = 0.011) indicate that mice with reduced *C9orf72* expression require more time to complete each stride, spend less time swinging the hindlimb through the air, and spend longer in contact with the ground. Additionally, C9-NRE significantly influenced Swing (*p* = 0.033) and Stance/Swing (*p* < 0.001), reiterating a shift in the balance of gait phases toward increased ground contact. Paw Drag and Propel also showed significant or near-significant genotype effects (*p*_PawDrag_ = 0.004; *p*_Propel_F_ = 0.041; *p*_Propel_M_ = 0.078), with Homo mice exhibiting larger paw drag areas and longer propulsion phases, consistent with increased effort during push-off and limb clearance.

**Figure 5:**
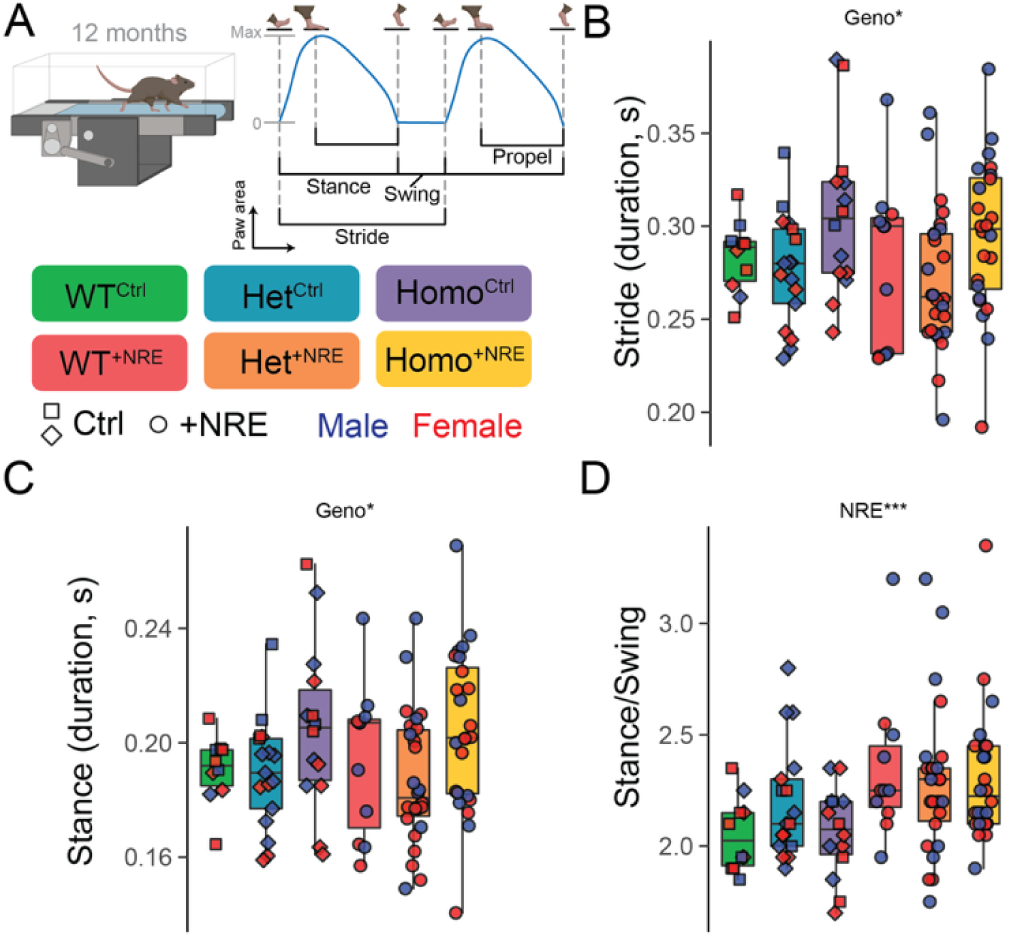
Subtle gait abnormalities present in ALS/FTD mice. (A) Schematic of paw area traces illustrating phases of gait. (B–C) *C9orf72* genotype affects hind paw stride duration (B) and stance time (C). (A) (D) Differences in hind paw stance-to-swing ratio are driven by C9-NRE. All plots are Tukey box plots. *n* = 3–16 per group.

In summary, these gait alterations indicate that both *C9orf72* genotype and C9-NRE expression are associated with measurable shifts in gait parameters at 12 months, characterized by increased time spent in ground contact and altered propulsion dynamics. Although these effects are subtle, they are consistent with early motor system dysfunction.

### Mild hyperactivity observed in the open field; anxiety remains unchanged

Given the widespread CNS expression achieved by intrathecal C9-NRE delivery (Figure 1), we also examined behavioral features relevant to the cognitive spectrum of ALS/FTD. Previous C9-NRE AAV studies assessing locomotion and exploratory behavior have reported variable outcomes, including hyperactivity, no detectable behavioral differences, or trends toward reduced general activity^29,43–46^. To determine where this intrathecal C9-NRE model falls along this behavioral spectrum, mice were subjected to open field testing at 10 months of age (Figure 6A). This assessment showed that C9-NRE mice spent comparable amounts of time in the center zone across genotypes and relative to Controls and that genotype did not significantly influence time in the center zone (Figure S5A), signifying no evidence of heightened, anxiety-like behavior in this model. In contrast, multiple measures of locomotion and exploratory activity were significantly altered. Total distance traveled, average speed, number of immobile episodes, perimeter zone speed, and perimeter zone distance traveled all showed significant main effects of genotype and C9-NRE expression (Figure 6B–C; Figure S5B). Additionally, total time spent mobile differed significantly across genotypes (Figure 6D).

**Figure 6:**
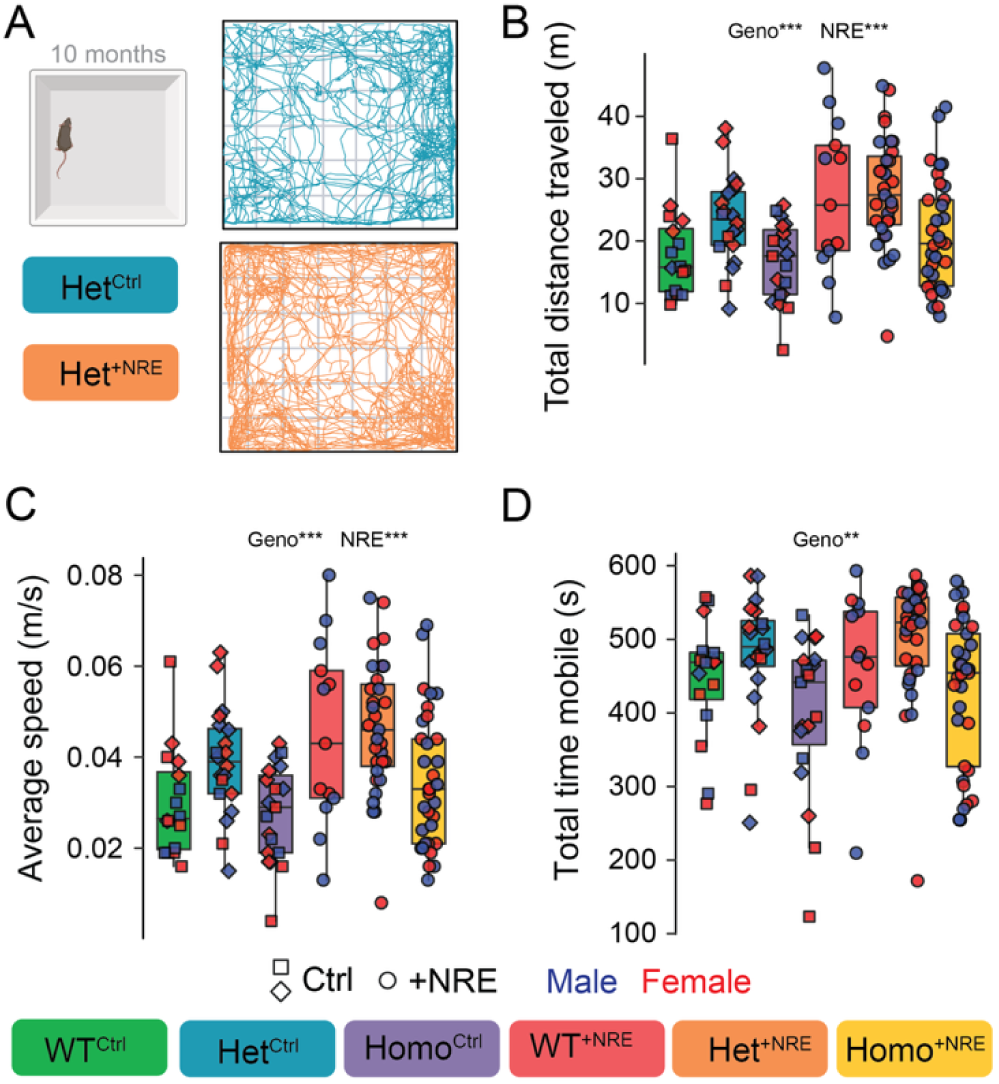
Open field assessment reveals mild hyperactivity. (A) Exported video traces of open field data showing the path representative mice traveled illustrate hyperactivity in C9-NRE mice and no evidence of anxiety-like behavior. (B–C) Total distance traveled (B) and average speed (C) are influenced by both *C9orf72* genotype and C9-NRE expression. (D) Total time spent mobile also differs by genotype. Box plots are Tukey box plots. *n* = 5–18 per group.

Together, these findings indicate increased locomotor activity in mice driven by both reduced *C9orf72* expression and the presence of C9-NRE expression.

### DPR pathology is robust throughout the cervical ventral horns in C9-NRE mice

Because DPRs are widely implicated as toxic contributors to ALS/FTD and consistently observed in preclinical C9-NRE models^29,43–46,63,64^, and given the motor and behavior phenotypes observed here, we assessed DPR pathology using IHC targeting poly-GP, poly-GR, and poly-GA in spinal cord tissue. Analyses were performed on the lower cervical spinal cord and focused on the ventral horns, as these regions contain motor neurons and interneurons involved in forelimb motor control^65^. At 12 months of age, robust DPR accumulation was detected in repeat-expressing mice. Poly-GP, poly-GR and poly-GA were readily observed throughout the ventral horns in diffuse and aggregated forms, whereas staining in Control animals was limited to background levels (Figure 7A–B). Quantitative analysis revealed a significant effect of C9-NRE expression on poly-GP, poly-GA (both *p* < 0.001), and poly-GR (*p* = 0.002) burden, as anticipated, with no significant differences across genotypes, indicating that DPR accumulation is driven by repeat expression. Longitudinal assessment across 6, 9, and 12 months showed a significant effect of time on poly-GA burden (Figure 7C). Poly-GP and poly-GR areas increased with positive trends over time, but these changes did not reach statistical significance, suggesting gradual accumulation that may require increased sampling to resolve temporal dynamics.

**Figure 7:**
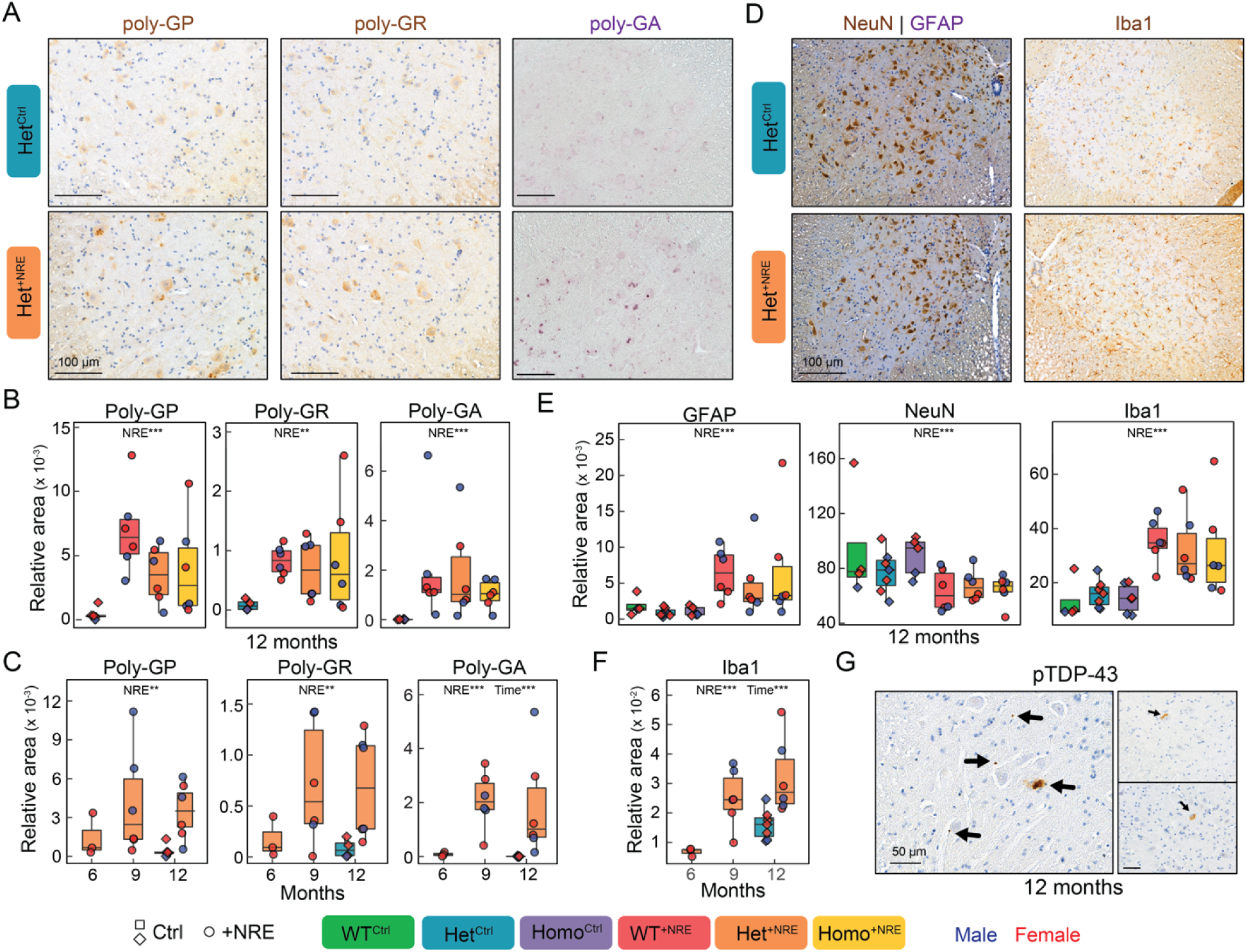
Hallmark C9-NRE ALS/FTD pathology detected throughout the cervical ventral horns. (A–B) Poly-GP, poly-GR, and poly-GA are detected in C9-NRE mice at levels significantly above background observed in Control animals. Representative images of lower cervical ventral horns (A) and quantification of DPR^+^ area relative to ventral horn area (B) at 12 months. DAB/brown: poly-GP (left), poly-GR (middle); VIP/purple: poly-GA (right); Hematoxylin counterstain. (C) Time significantly affects poly-GA area. Although poly-GP and poly-GR areas show a positive trend from 6 to 12 months, these changes do not reach statistical significance. (D–E) C9-NRE mice show increased GFAP (E, middle) and Iba1 (E, right) areas, indicating astrogliosis and microgliosis, and decreased NeuN area (E, left), signifying smaller and/or fewer neurons, compared to Controls. Representative images of lower cervical ventral horns stained for NeuN, GFAP, and Hematoxylin (D, left panels) and Iba1 and Hematoxylin (D, right panels) at 12 months. DAB/brown: NeuN and Iba1; VIP/purple: GFAP (D, left only). (F) Iba1 area increases over time; 9- and 12-month-old Het^+NRE^ mice show significantly more microgliosis than 6-month-old Het^+NRE^ mice. (G) pTDP-43 pathology, including small puncta, larger aggregates, and diffuse, cytoplasmic staining is present in C9-NRE tissue. Representative images of pTDP-43 pathology (arrows) in the grey matter of lower cervical tissue. DAB/brown: pTDP-43 and Hematoxylin. All box plots are Tukey box plots.

Together, these findings indicate that intrathecal injection of C9-NRE is sufficient to elicit substantial DPR accumulation in pathophysiologically relevant regions of the cervical spinal cord, regardless of *C9orf72* genotype.

### C9-NRE expression reduces NeuN-positive area and promotes progressive neuroinflammatory pathology

Progressive motor neuron degeneration and glial activation are central pathological features of ALS/FTD, with both patient tissue^30–39^ and preclinical models^29,40–46^ exhibiting neuronal loss, astrogliosis, and microgliosis. In light of this and the pathological and motor phenotypes observed above, we assessed whether these hallmarks are recapitulated in this model by performing IHC analyses in the ventral horns of the lower cervical spinal cord. We assessed neuronal area using NeuN, astrocytic reactivity using GFAP, and microglial burden using Iba1. At 12 months, C9-NRE expression was associated with significantly reduced NeuN area in ventral horn regions compared with non-repeat-expressing Controls (*p* < 0.001; Figure 7D–E), an effect confirmed by pairwise contrasts within each genotype. Additionally, GFAP and Iba1 areas were significantly increased in the ventral horns of 12-month-old C9-NRE mice compared with Controls (both *p* < 0.001; Figure 7D–E). Pairwise comparisons in both of these measures reflected this, showing increased astrocyte and microglia area in C9-NRE mice relative to genotype-matched Controls. *C9orf72* genotype did not significantly influence NeuN, GFAP, or Iba1 area.

Given the robust increase in microglial burden, we next examined whether Iba1 pathology evolved over time. Longitudinal analysis of Iba1 further revealed progressive microgliosis, with a significant effect of time on Iba1 area (*p* < 0.001), with microglial burden significantly lower at 6 months than at both 9 and 12 months (both *p* < 0.001; Figure 7F). Though still trending upwards, there was no significant difference between Iba1 area at 9 and 12 months (*p* = 0.198), suggesting either an early influx that progresses to a chronic state of neuroinflammation, maintained through later disease stages or again, the need for increased sampling to resolve temporal dynamics.

Collectively, these data demonstrate that intrathecal C9-NRE expression drives progressive spinal cord pathology characterized by astrocytic activation, age-dependent microgliosis, and reduced NeuN-positive area.

### Sparse pTDP-43 pathology is observed in C9-NRE grey matter

TDP-43 pathology, observed in the majority of ALS/FTD cases, involves both gain- and loss-of-function mechanisms and features cytoplasmic mislocalization of abnormally phosphorylated TDP-43 (pTDP-43), leading to toxic aggregate formation and neurodegeneration^66,67^. Given the central role of TDP-43 pathology in ALS/FTD and the mixed reports of TDP-43 pathology across existing mouse models, we performed IHC analyses to determine whether disease-associated pTDP-43 is detectable in the spinal cord of this gain- and loss-of-function model. We observed sparse pTDP-43 pathology in the gray matter of cervical spinal cord sections from 12-month-old C9-NRE mice, appearing as small puncta, larger aggregates, and diffuse cytoplasmic staining (Figure 7G). pTDP-43-positive profiles were infrequent and each was, expectedly, limited to one or two sections, precluding robust quantitative analysis. Scattered dot-like pTDP-43 staining was also detected in some 9-month tissues (Figure S6), potentially reflecting early or pre-aggregate pathology. Though sparse, the detection of pTDP-43 pathology demonstrates this model’s potential to recapitulate hallmark features of ALS/FTD.

### C9-NRE expression level variability and cross-domain correlation analyses support a mild, disease-relevant ALS/FTD model

To more rigorously account for variability in C9-NRE expression levels achieved by this intrathecal AAV approach and its potential relationship to phenotypic and pathological outcomes, repeat expression was quantified in the upper cervical spinal cord across all animals that completed 12-month testing and those used for IHC analyses. Median C9-NRE expression was 143.74-fold relative to GAPDH, with a mean expression of 412.80-fold relative to GAPDH (Figure 8A). No differences in cervical C9-NRE expression were observed across genotypes or between sexes (Table S1), indicating that variability in repeat expression was not systematically associated with these biological variables.

**Figure 8:**
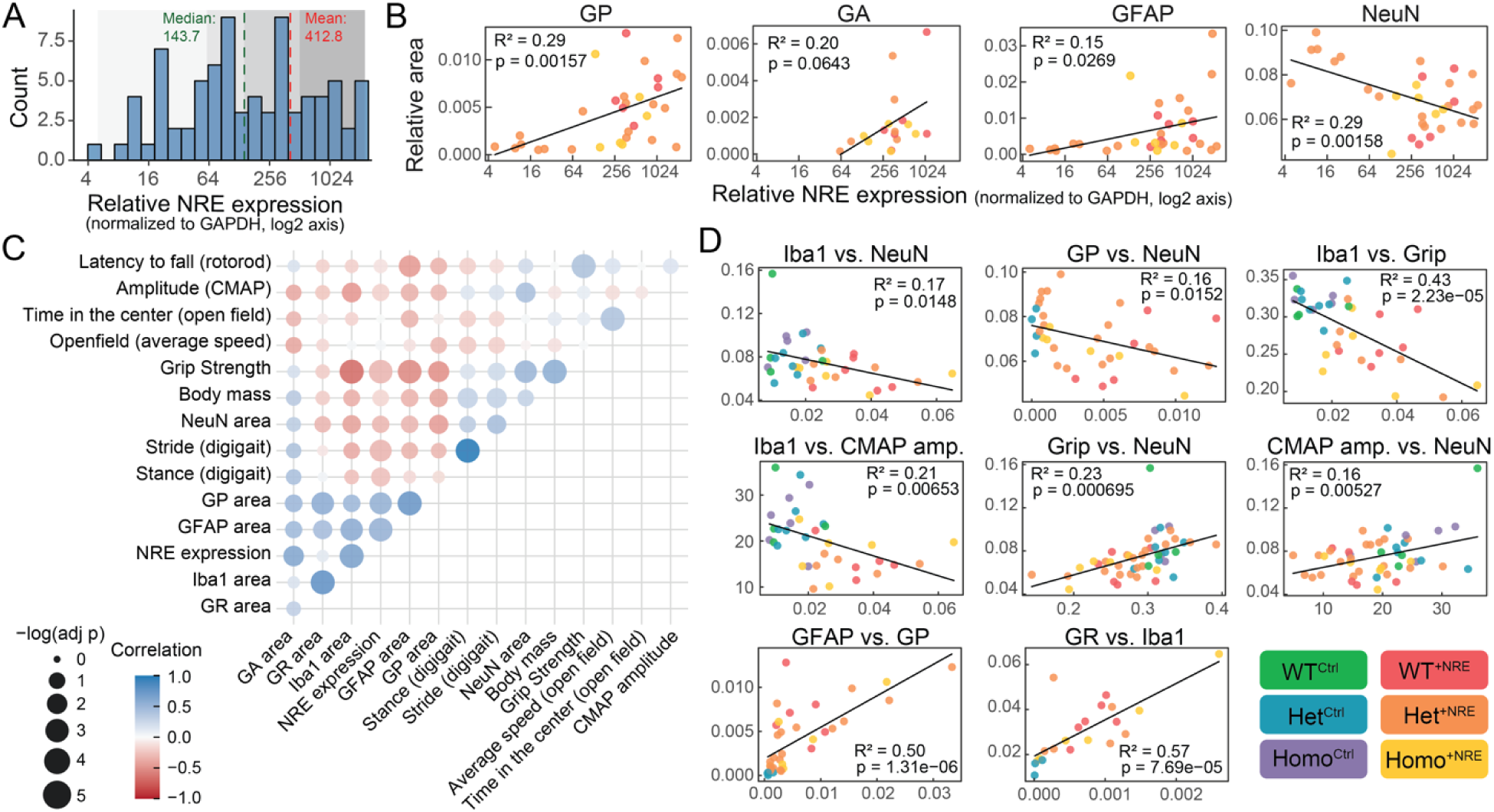
Correlations among variables support pathophysiological links. (A) Histogram of C9-NRE expression levels in upper cervical spinal cord, normalized to GAPDH expression; shading indicates quartiles. (B) At 12 months, linear regression shows C9-NRE expression level correlates with poly-GP, poly-GA, GFAP, and NeuN areas. (C) Pearson correlation analysis at the last experimental time points reveals relationships among representative measures of motor performance, behavior, spinal cord pathology, and C9-NRE expression levels; color indicates strength and direction of correlation, and dot size reflects significance. (D) Additional regressions of variables shown in (C) illustrate disease-relevant links between pathology and phenotypes in this model. Variables are plotted as listed (x-axis, y-axis).

Given this repeat expression distribution, linear regression analyses using the least-squares method were applied to examine whether variability in C9-NRE expression is associated with differences in pathological outcomes. These analyses showed that poly-GP, poly-GA, GFAP, and NeuN areas significantly or near-significantly correlated with C9-NRE expression levels, with higher repeat expression associating with increased poly-GP and poly-GA burden, greater astrocytic reactivity (i.e. astrogliosis), and reduced NeuN area (i.e., decreased neuronal number and/or size) (Figure 8B). In contrast, C9-NRE expression level did not significantly correlate with poly-GR area or Iba1 area (Figure S7A), suggesting that variability in repeat expression may be more closely linked to select pathological features than to others.

Relationships across functional and pathological domains were then assessed to determine how molecular, cellular, and behavioral measures covary in this intrathecal C9-NRE model. Pearson correlation analysis incorporating representative measures of motor performance, behavior, spinal cord pathology, and C9-NRE expression levels revealed coherent cross-domain relationships, with biologically related measures clustering together (Figure 8C). Pathological markers exhibited strong internal coherence, with DPRs, GFAP, and Iba1 positively correlating with one another, inversely correlating with NeuN area, and showing largely similar association patterns with functional measures. Minor exceptions included poly-GR, which showed weak correlations with gait parameters, and poly-GA, which showed slight deviations in correlation patterns with some functional measures. Gait parameters exhibited modest correlations with both pathological and motor strength measures, supporting gait as a sensitive but multifactorial readout influenced by neuromuscular and inflammatory states. Motor strength measures clustered together, and correlated positively with NeuN and negatively with pathological markers. Rotarod showed weaker and partially distinct associations, consistent with its greater dependence on coordination. Open field measures largely segregated from motor and pathological domains, indicating hyperactivity represents a partially independent behavioral phenotype.

Guided by these domain-level patterns, we performed linear regression to further probe pairwise relationships among phenotypic measures showing significant correlations (*p* < 0.05, Figure 8D; Figure S7B). Several associations were consistent with established disease mechanisms, including positive correlations between poly-GP and poly-GR and between Iba1 and GFAP, reflecting concordance among markers of RAN translation and neuroinflammation, respectively. Inverse correlations between Iba1 and NeuN and between poly-GP and NeuN linked inflammatory and DPR pathology to reduced neuronal area. Motor deficits also correlated with inflammatory burden, with Iba1 associating with grip strength and CMAP amplitude and GFAP associating with rotarod and grip strength. Reduced NeuN area similarly correlated with decreased grip strength and CMAP amplitude, connecting neuronal decline to functional deficits. The lower cervical ventral horns examined here govern forelimb motor control, reinforcing the link between spinal pathology and function, and given the widespread spinal C9-NRE expression, similar coupling likely extends to regions governing hindlimb function. Together, these relationships demonstrate that engagement of pathophysiologically relevant regions translates to measurable, disease-relevant motor phenotypes.

## DISCUSSION

Preclinical mouse models of C9-NRE ALS/FTD incompletely capture motor phenotypes and fail to robustly engage the spinal cord, despite lower motor neuron degeneration and motor dysfunction being defining features of disease. To address these gaps, we combined neonatal intrathecal delivery of a pathogenic C9-NRE AAV with graded loss of endogenous *C9orf72*, targeting both interacting molecular mechanisms and disease-relevant motor regions. Using this approach, we demonstrate that integrating both gain- and loss-of-function mechanisms with CNS-wide AAV engagement produces convergent motor, behavioral, and pathological features consistent with a mild but biologically meaningful ALS/FTD phenotype.

Intrathecal AAV2/9 delivery recapitulated brain expression patterns reported in prior ICV-based C9-NRE studies^29,45^ while also achieving robust repeat expression throughout the spinal cord. In the context of reduced *C9orf72* function, C9-NRE expression emerged as the primary driver of progressive muscle weakness (grip strength and electrophysiology), whereas impairments in motor coordination (rotarod) were more closely aligned with *C9orf72* genotype. While mice did not exhibit overt anxiety-like behavior, they displayed hyperactivity in the open field and subtle, consistent gait abnormalities.

This model reveals a dissociation between strength- and coordination-dependent phenotypes, reflecting differential effects of repeat toxicity and *C9orf72* loss on motor circuits. Concurrent reductions in CMAP amplitude and area are consistent with motor axon degeneration and/or muscle atrophy rather than altered conduction or neuromuscular junction dysfunction^68^. These electrophysiological findings align with grip strength deficits and indicate that C9-NRE expression drives motor unit degeneration, with genotype-dependent effects emerging later, further exacerbating muscle weakness. In contrast, genotype-associated rotarod deficits parallel those reported in the prior combinatorial ICV model^29^, where expression and pathology implicated brain regions involved in motor control.

Although gait abnormalities were subtle, they resemble early motor changes reported in other neurodegenerative models. Longer stride and stance times mark early gait deficits in SOD1 mice^61^, and a preclinical Huntington’s disease model shows an increased proportion of stride spent in stance^69^. Increased propulsion time and paw drag also occur in SOD1 mice as early indicators of motor impairment^62^. These alterations in gait are consistent with early motor system compromise and reflect the multifactorial nature of gait, which integrates muscle strength, coordination, and balance. The partial overlap between genotype- and repeat-driven effects in this model is therefore not unexpected and likely reflects integration across multiple motor control pathways affected in ALS/FTD.

In this model, hyperactive behavior emerged alongside a lack of anxiety-related phenotypes and persisted within the perimeter zone, typically associated with safety, supporting hyperactivity as a primary behavioral feature rather than an anxiety-driven response. These findings align with multiple C9-NRE^43,44,46,70^ and ALS/FTD^71–74^ preclinical models and mirror behavioral disinhibition reported in subsets of FTD patients^75,76^.

Pathologically, intrathecal C9-NRE expression in *C9orf72* loss-of-function mice induced DPR accumulation, reduced NeuN-positive area, activated astrocytes and microglia, and produced sparse pTDP-43 pathology– hallmarks observed in patient tissue and preclinical ALS/FTD models^29,43–46^. Importantly, cross-domain correlation analyses confirmed relationships linking repeat expression, spinal pathology, and motor dysfunction, reinforcing the model’s disease relevance despite its modest phenotypic severity. These analyses revealed additional intriguing relationships, including a divergence between DPR species, with poly-GP associating more strongly with astrocytic activation (GFAP) and poly-GR associating more modestly with GFAP but strongly with microglial activation (Iba1) (Figure 8D; Figure S7B), suggesting distinct relationships between specific DPR species and inflammatory subtypes.

Collectively, these findings demonstrate that CNS-wide C9-NRE expression, including the spinal cord, in the context of reduced *C9orf72* function produces coordinated molecular, cellular, and functional abnormalities that recapitulate key aspects of ALS/FTD. While the overall phenotype is mild, the coherence across motor, behavioral, and pathological domains supports the model’s utility for interrogating disease mechanisms, temporal relationships between pathology and function, and therapeutic interventions targeting motor and spinal cord-associated pathology.

### Limitations and Future Directions

Although repeat expression was robust and widespread, resulting phenotypes were modest and most pronounced at late time points, suggesting that disease severity in this model may be constrained by repeat expression levels, age, and/or the balance between gain- and loss-of-function mechanisms. Higher AAV doses may amplify reported phenotypes but must be approached cautiously, as they can induce inflammation or toxicity^77–80^, and consideration of serotype, promoter, and route of delivery would be necessary to mitigate off-target effects^50,81–83^. A number of phenotypes were time-dependent, and possible evidence of gain- and loss-of-function interactions emerged only at later time points, indicating that aging the model beyond 12 months may further enhance phenotypic divergence, as well as clarify synergistic relationships between pathogenic mechanisms.

Un-injected and dye-injected controls account for *C9orf72* loss-of-function effects and neonatal intrathecal injection stress, respectively; however, inclusion of a non-disease-causative (G_4_C_2_)_2_ AAV would additionally control for potential AAV-related effects. A prior study^45^ using a comparable C9-NRE AAV reported no behavioral or pathological differences between sham and (G_4_C_2_)_2_ Controls and minimal divergence in their comparisons to (G_4_C_2_)_66_ animals, indicating equivalence of sham and (G_4_C_2_)_2_ Controls. Another study using similar AAV dosing reported no motor, neurodegeneration, or transcriptome effects in AAV-GFP Controls^54^. Together, these findings suggest that the phenotypes observed here are unlikely to reflect nonspecific AAV effects; nonetheless, inclusion of such controls would strengthen future studies.

Pathological analyses were necessarily limited by IHC sampling constraints. Sparse pTDP-43 pathology was detected, but its infrequency precluded robust quantification. Sequential sectioning, whole-tissue analyses, or complementary biochemical approaches may be required to fully characterize TDP-43 pathology and its temporal evolution in this model. Additionally, as mentioned above, more extensive IHC sampling might further clarify other temporal pathological dynamics in this model.

Finally, this study focused pathologically on the cervical motor region; however, given that ALS/FTD is a multisystem disease, future work incorporating lumbar/lower limb spinal regions, as well as detailed cortical, cerebellar, brainstem, muscular, and circuit-level analyses, will be important for fully defining how distinct regional and cellular mechanisms interact to drive disease progression. Cell-type-specific manipulations will also be valuable to clarify selective vulnerability and parse the contributions of pathophysiological processes, including neuroinflammation, to disease.

In summary, despite these limitations, this intrathecal C9-NRE model provides a biologically coherent and disease-relevant framework for studying ALS/FTD. By linking repeat expression, spinal pathology, and functional decline, it offers a platform for mechanistic dissection and therapeutic evaluation, particularly for interventions targeting early or progressive motor dysfunction.

## RESOURCE AVAILABILITY

Requests for further information and resources should be directed to the lead contact, Aaron Haeusler.

This study did not generate new unique reagents.

Any additional information required to reanalyze the data reported in this paper is available from the lead contact upon request.

## ACKNOWEDEGMENTS

We thank members of the Jefferson Weinberg ALS Center for valuable feedback throughout the study, Dr. Leonard Petrucelli and colleagues for kindly providing AAV constructs and poly-GA antibody, Mo Singer for establishing DigiGait processing workflows, and Matt Davis for technical assistance with qPCR. This work utilized the Translational Pathology Shared Resource of the Sidney Kimmel Comprehensive Cancer Center, Thomas Jefferson University (supported by NCI grant 5P30CA056036). Figures include schematics created with BioRender.com. Funding was provided by RF1NS114128 and R01NS114128 to ARH; R01NS109150 to BKJ; and the Farber Family Foundation and Aldrich Foundation to the Jefferson Weinberg ALS Center.

## AUTHOR CONTRIBUTIONS

KAR and ARH conceptualized the project. BKJ performed qPCR. SCA completed poly-GA staining, imaging, and analyses. JDR oversaw poly-GA analyses. KAR developed the methodology and completed all other experiments. AAS executed video processing. KAR, AAS, and BKJ pre-processed IHC images. KAR performed mixed model analyses. ARH conducted correlation analyses, R programming, and machine learning-based feature predictions, and oversaw data analysis. KAR led manuscript writing, with ARH. KAR and ARH developed and assembled the figures. MDB advised on IHC staining. BKJ, JDR, and DT provided manuscript feedback. All authors approved the final manuscript.

## DECLARATION OF GENERATIVE AI AND AI-ASSISTED TECHNOLOGIES IN THE WRITING PROCESS

During the preparation of this work, the authors used Large Language Models, such as ChatGPT and Grammarly, to improve grammar and clarity. After using this tool/service, the authors reviewed and edited as needed and take full responsibility for the content of the publication.

