## SUPPLEMENTAL FIGURES AND FIGURE LEGENDS for "Intrathecal (G_4_C_2_)_149_ delivery in C9orf72-deficient mice yields mild motor dysfunction and ALS/FTD pathological hallmarks"

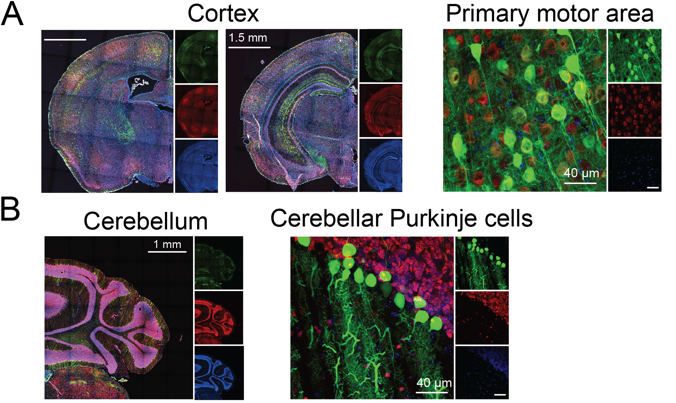

**Figure S1: Intrathecal AAV delivery attains GFP reporter expression throughout the brain.**

(A–B) Intrathecal injections of AAV2/9 reporter show GFP expression throughout the brain, including disease-relevant regions, as similarly reported in previous ICV AAV injection studies^29,43,45^. The primary motor area, hippocampus, and neurons within the primary motor area exhibit robust GFP signal (A, top row). GFP expression is also prolific throughout the cerebellum, specifically in Purkinje cells (B, bottom row). Tissue was harvested four weeks post-injection. Green: GFP; Red: NeuN; Blue: Hoechst.

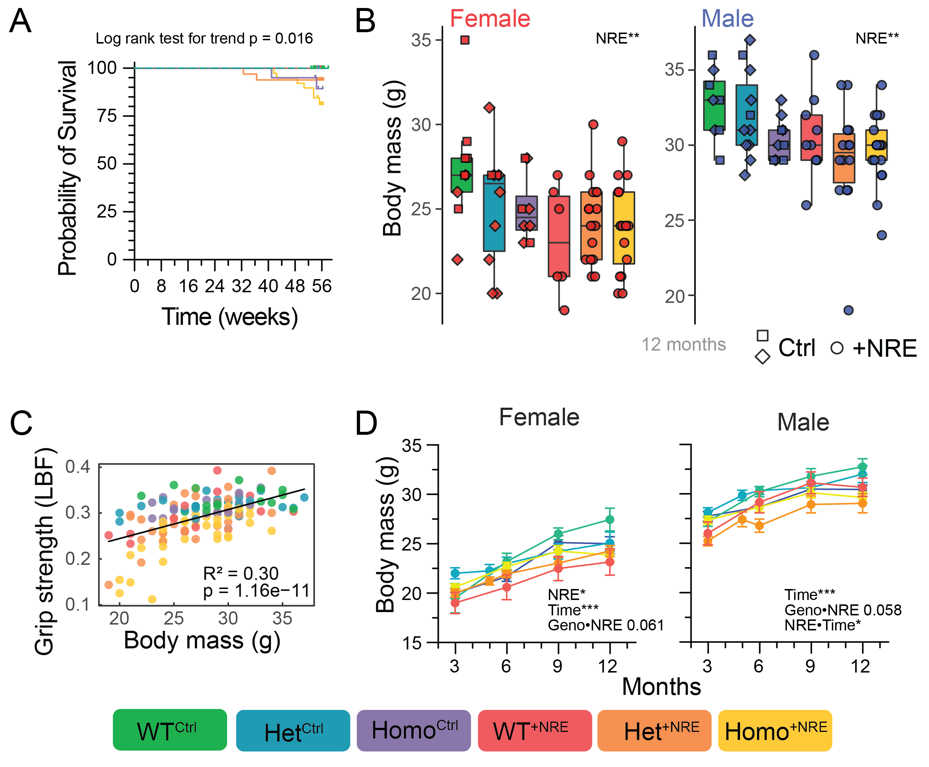

**Figure S2: Mice display similar survival probability across cohorts, while body mass varies by sex and C9-NRE expression.**

(A) At 12 months of age, survival did not differ significantly across cohorts by log-rank (Mantel–Cox) or Gehan–Breslow–Wilcoxon tests, but showed a significant trend by the log-rank test for trend (ordered by WT^Ctrl^, Het^Ctrl^, Homo^Ctrl^, WT^+NRE^, Het^+NRE^, Homo^+NRE^).

(B) At 12 months of age, male mice weighed more than female mice and C9-NRE mice weighed less than Control mice (both *p* < 0.001). Differences between C9-NRE and Control body mass persist after stratification by sex.

(C) Simple linear regression using the least squares method revealed significant correlation between grip strength and body mass; however, this relationship was driven, at least in part, by differences between Ctrl and C9-NRE animals, as evidenced by the clustering of repeat-containing mice toward the lower left quadrant.

(D) Longitudinal analysis of body mass showed persistent effects of the repeat expansion, with C9-NRE and several of its interactions identified as significant drivers. Overall, C9-NRE mice exhibited lower body masses than Control mice.

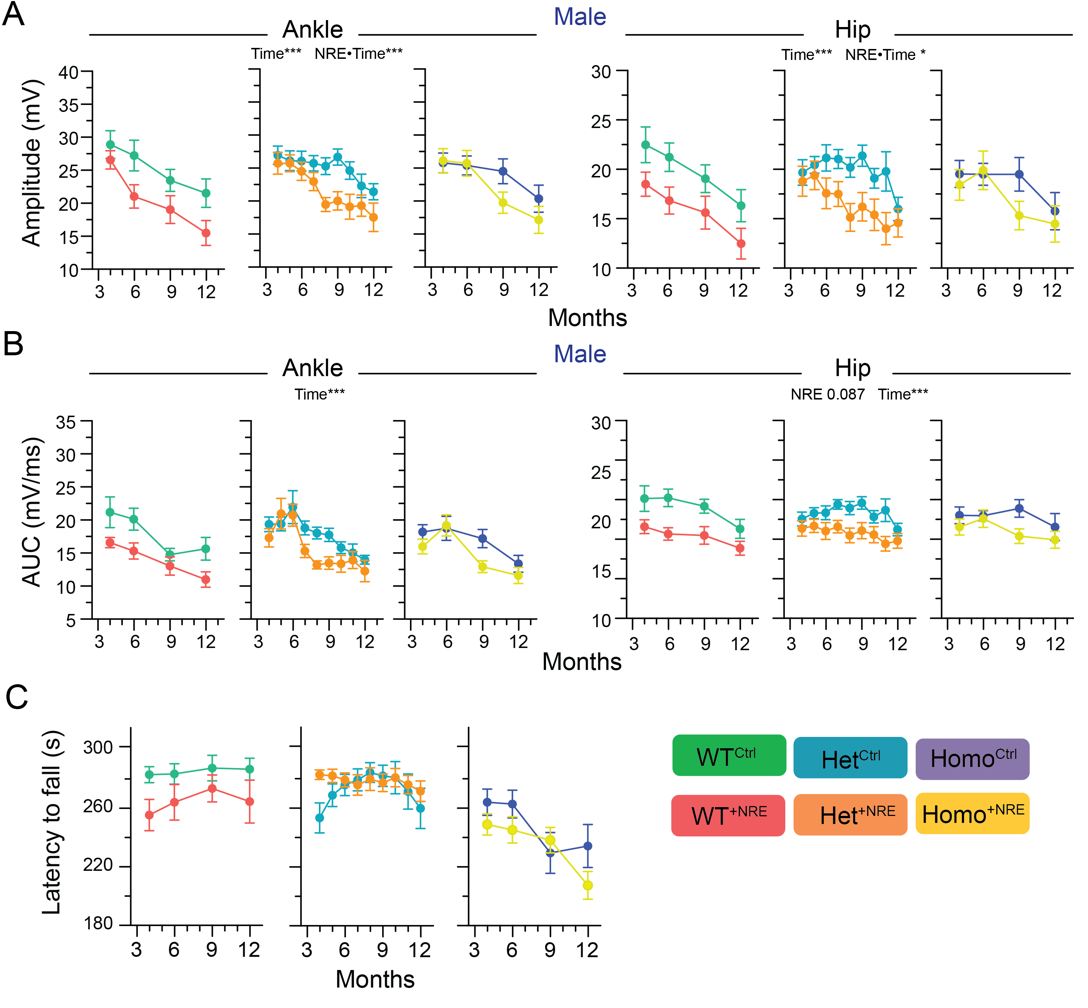

**Figure S3: Male mice recapitulate C9-NRE-dependent electrophysiological phenotypes and rotarod data graphed by genotype emphasizes *C9orf72*-dependent motor coordination.**

(A–B) Similar to female mice, male mice show C9-NRE-dependent motor deficits in measures of both CMAP amplitude and area. C9-NRE mice have lower CMAP amplitudes (A) and decreased CMAP areas (B) compared to Control mice, suggesting motor strength deficits are repeat expansion-dependent.

(C) Rotarod data over time, now paired by genotype, reiterates reduced performance is driven by *C9orf72* genotype. Regardless of the presence of the expanded repeat, Homo mice produce shorter latencies to fall compared to their Het and WT counterparts. Line graphs show mean ± SEM.

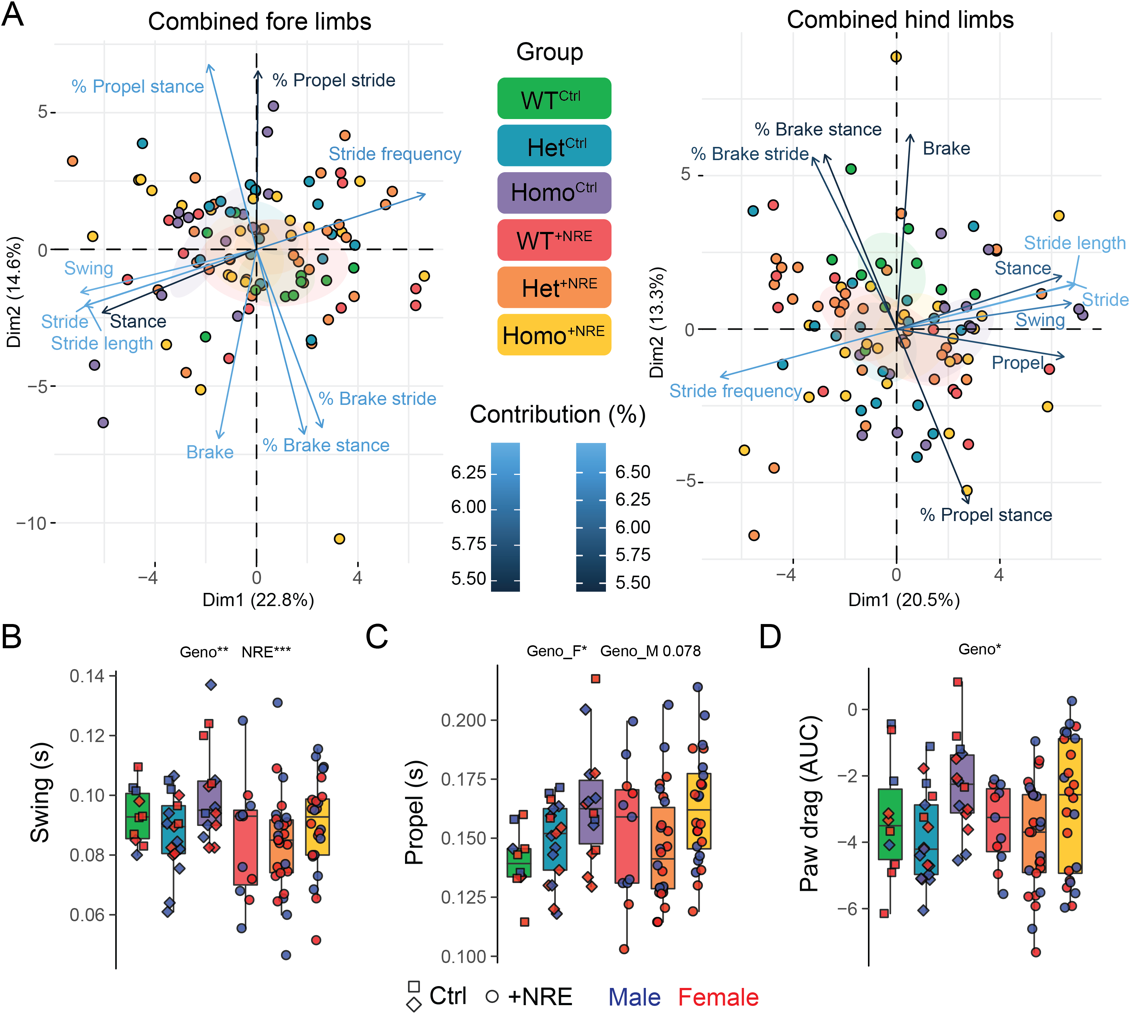

**Figure S4: Genotype-dependent alterations in hind paw gait.**

(A) PCA plots for both fore paws and hind paws showing the top ten contributing variables. Hind paws are the focus of subsequent analysis.

(B) Swing, the time that hind paws spend not in contact with the ground, shows differences across both C9-NRE and genotype.

(C–D) Propel, the time hind paws are pushing off the ground, and Paw drag, the area under the curve during propulsion, are both significantly influenced by genotype.

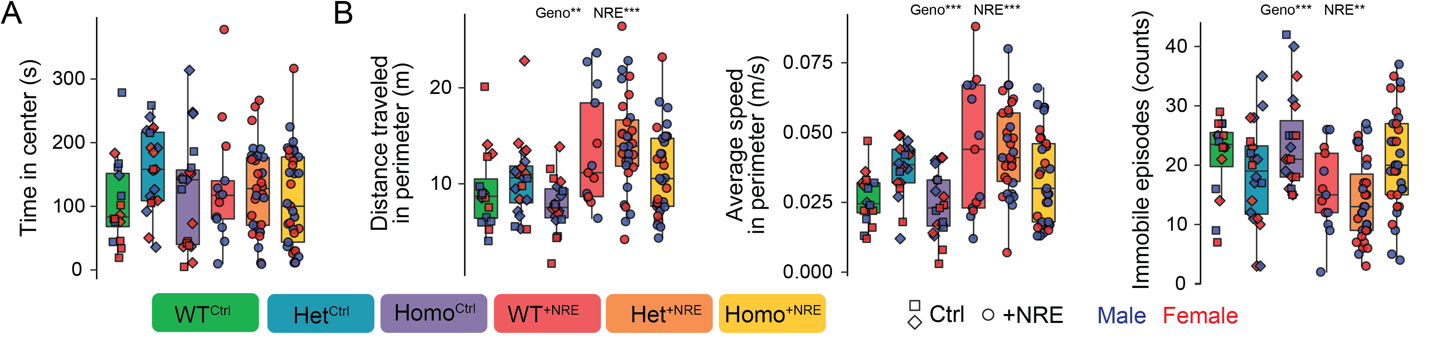

**Figure S5: Open Field tests show no evidence of anxiety, but hyperactivity observed, persisting in the perimeter zone.**

(A) Mice spend equivalent amounts of time in the center zone of the open field, not influenced by genotype or C9-NRE.

(B) A hyperactive phenotype is reiterated by increased distance traveled and average speed phenotypes persisting when focusing analysis on just the perimeter zone (left and middle panels), as well as fewer instances of immobility overall (right panel).

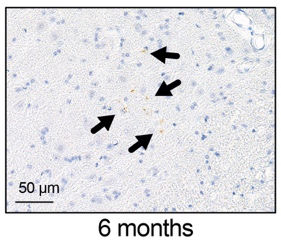

**Figure S6: Early pTDP-43 pathology observed in cervical spinal tissue at 9 months.**

Ventral horns from the lower cervical spinal cord, harvested at 9 months of age and stained for pTDP-43, show a scattered, dot-like staining pattern, potentially indicating either early or pre-aggregate pathology. DAB/brown: pTDP-43.

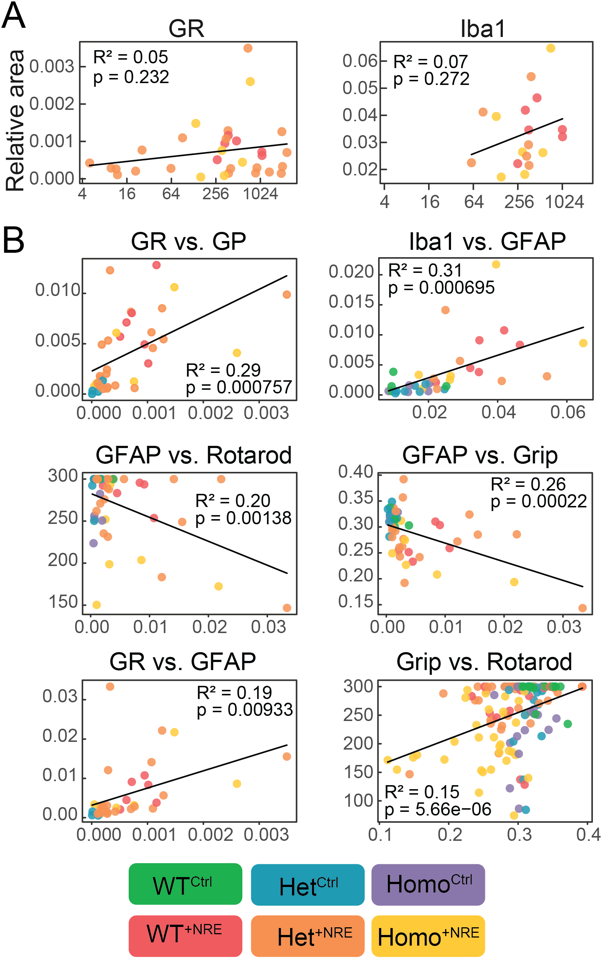

**Figure S7: Correlations provide insight into connections among C9-NRE expression level, pathology, and functional readouts.**

(A) Simple linear regression using the least squares method shows C9-NRE expression level does not significantly correlate with either poly-GR or Iba1 at the 12-month time point.

(B) Additional regressions plot various relationships between experimental readouts highlighting correlations between similar variables (first row), relationships between pathology and function (second row), as well as shedding light on some potentially interesting avenues for further study (third row).

**Table S1: Mixed model *p* values for experimental measures.**

Values reflect genotype, C9-NRE expression, sex, time, and interaction effects. See *Methods* for details regarding analysis workflow.
